# A virtual host model of *Mycobacterium tuberculosis* infection identifies early immune events as predictive of infection outcomes

**DOI:** 10.1101/2021.11.08.467840

**Authors:** Louis R. Joslyn, Jennifer J. Linderman, Denise E. Kirschner

## Abstract

Tuberculosis (TB), caused by infection with *Mycobacterium tuberculosis* (Mtb), is one of the world’s deadliest infectious diseases and remains a significant global health burden. TB disease and pathology can present clinically across a spectrum of outcomes, ranging from total sterilization of infection to active disease. Much remains unknown about the biology that drives an individual towards various clinical outcomes as it is challenging to experimentally address specific mechanisms driving clinical outcomes. Furthermore, it is unknown whether numbers of immune cells in the blood accurately reflect ongoing events during infection within human lungs. Herein, we utilize a systems biology approach by developing a whole-host model of the immune response to Mtb across multiple physiologic and time scales. This model, called *HostSim*, tracks events at the cellular, granuloma, organ, and host scale and represents the first whole-host, multi-scale model of the immune response following Mtb infection. We show that this model can capture various aspects of human and non-human primate TB disease and predict that biomarkers in the blood may only faithfully represent events in the lung at early time points after infection. We posit that *HostSim*, as a first step toward personalized digital twins in TB research, offers a powerful computational tool that can be used in concert with experimental approaches to understand and predict events about various aspects of TB disease and therapeutics.

## Introduction

Even during the COVID-19 pandemic, tuberculosis (TB) continues to be a global threat. Approximately 25% of the world is infected with *Mycobacterium tuberculosis* (Mtb) and 5-10% of those currently infected will progress to develop symptomatic clinical disease (1). TB patients are often classified as having latent tuberculosis (LTBI) or active TB. LTBI is an asymptomatic state of infection with typically low levels of Mtb present. Active TB cases exhibit clinical symptoms including fever, weight loss, night sweats, and coughing typically with high levels of Mtb present. While patients are categorized within these binary states, recent work has shown that TB manifests as a spectrum of clinical and infection outcomes within humans and non-human primates (NHPs) (2–5). LTBI individuals can undergo reactivation events and therefore act as a potential reservoir for disease transmission (6,7). Much remains unknown about the biology that drives clinical outcomes in TB (i.e., latent or active) for each individual patient. It is critical to understand events that lead to latent or active TB in order to develop effective vaccines and host-directed therapies.

The hallmark of TB is the formation of lung granulomas, which are organized immune structures that surround Mtb and Mtb-infected cells within lungs of infected hosts (8). NHP data have shown that a single mycobacterium is sufficient to begin the formation of a granuloma and that each granuloma has a unique trajectory (9,10). Granulomas are composed of bacteria and various immune cells, such as macrophages and T cells (primarily CD4+ and CD8+ T cells, although other unconventional T cell phenotypes are also present, reviewed in (11)). Other cells such as neutrophils, fibroblasts and dendritic cells are also present. T cells have well-known critical functions against Mtb (12–14), but unlike other infections, T cells are slow to enter the site of infection within lungs, arriving approximately one month after primary infection (15). Lung-draining lymph nodes (LN) serve as sites for initiating and generating an adaptive immune response against most pulmonary infections, including Mtb. However, delays in LN T-cell priming, activation, and trafficking through blood to lungs is characteristic of the adaptive immune response in Mtb (16,17) and is thought to be key in allowing Mtb to establish infection within lungs (15). The delay is thought to arise from slowly growing mycobacteria in the lungs, delaying the signals for adaptive immunity (18).

While studies at the granuloma scale have elucidated important features about how individual granulomas control infection, it is difficult to experimentally identify immune mechanisms within lung granulomas and LNs that drive clinical outcomes of TB at a whole-host scale. Mediators such as CD4+ T cells, CD8+ T cells and TNFα are important in controlling established Mtb infection (12,19,20). NHP studies have shown that active TB individuals harbor significantly more bacteria than LTBI individuals (21) but these studies have been unable to relate individual granuloma outcomes to whole-host clinical outcomes, in part because the fate of individual granulomas vary within a single host (9).

Data from sites of infection (lung granulomas) in humans are generally unavailable. Consequently, it is not known whether numbers of immune cells in the blood reflect ongoing events during infection within human lungs (22). This has limited the ability to use blood as a predictive measure for infection progression or diagnosis. However, recent association studies suggest ratios of antigen-specific CD4+ and CD8+ T cells within the blood of Mtb-infected hosts may help delineate LTBI from active TB (23,24). Conversely, NHP studies have shown that T-cell responses in the blood do not consistently reflect T-cell responses in granulomas (25,26).

Mathematical and computational modeling offer complementary approaches to experimental studies. Models have the power to simultaneously track multiple immune cell populations across multiple compartments, explore mechanisms of action related to immunological phenomenon, and predict timing of major immune events. In TB (27), modeling has been used to explore bacterial behavior in relation to the granuloma environment (28), drug-dynamics within granulomas (29,30) and immune cell interactions and cytokines within a lung model (31–34). Additionally, pseudo whole-host models have been developed to begin to investigate biomarkers in TB (26) and drug dynamics across a host (35). Mathematical and computational modeling is a unique tool that could serve to bridge events occurring within a host to whole-host level TB outcomes (i.e. LTBI vs active TB).

Here we develop a novel whole-host scale modeling framework that captures key elements of the immune response to Mtb within three physiological compartments - LNs, blood and lungs of infected individuals. Beginning with our whole lung framework originally called *MultiGran*, each granuloma is formulated as an individual ‘agent’ in an agent-based model that contains a sub-model tracking immune cells, cytokines, and bacterial populations for each granuloma (36). We extend this framework to capture dynamics of a whole host by linking it with a two-compartment model representing immune cell dynamics occurring within LNs and blood (37,38). Together, this new model platform, called *HostSim*, represents a whole-host framework for tracking Mtb infection dynamics within a single host across long time scales (days to months to years). We calibrate and validate the model using multiple datasets from published NHP studies.

Once developed, we use *HostSim* to answer two outstanding questions surrounding whole-host outcomes in TB: 1) what are mechanisms within a host that drive clinical outcomes in TB at the whole-host scale? 2) is there a relationship between blood immune cell counts and clinical outcomes at the whole-host scale? We use *HostSim*, the first whole host multi-scale model of Mtb infection, to relate immune responses in the blood to the sites of infection within lungs. Additionally, we utilize sensitivity analysis to predict factors that lead to clinical outcomes of TB.

## Methods

### HostSim model overview

Our novel multi-scale whole-host scale model, *HostSim*, tracks Mtb infection across three separate physiological compartments (Figure 1). We describe the formation, function and dissemination of multiple granulomas that represent distinct sites of infection developing within a whole-lung model. We additionally describe the initiation of adaptive immunity within a LN compartment after receiving signals from antigen presenting cells migrating from lungs. Finally, we track immune cell counts within a blood compartment that acts as a bridge between LN to lungs. *HostSim* uses rule-based agent placement, employs parameter randomization, solves non-linear systems of ordinary differential equations (ODEs), performs post-processing agent groupings, and utilizes rule-based linking between scales to perform *in silico* simulations of a single host.

**Figure 1:**
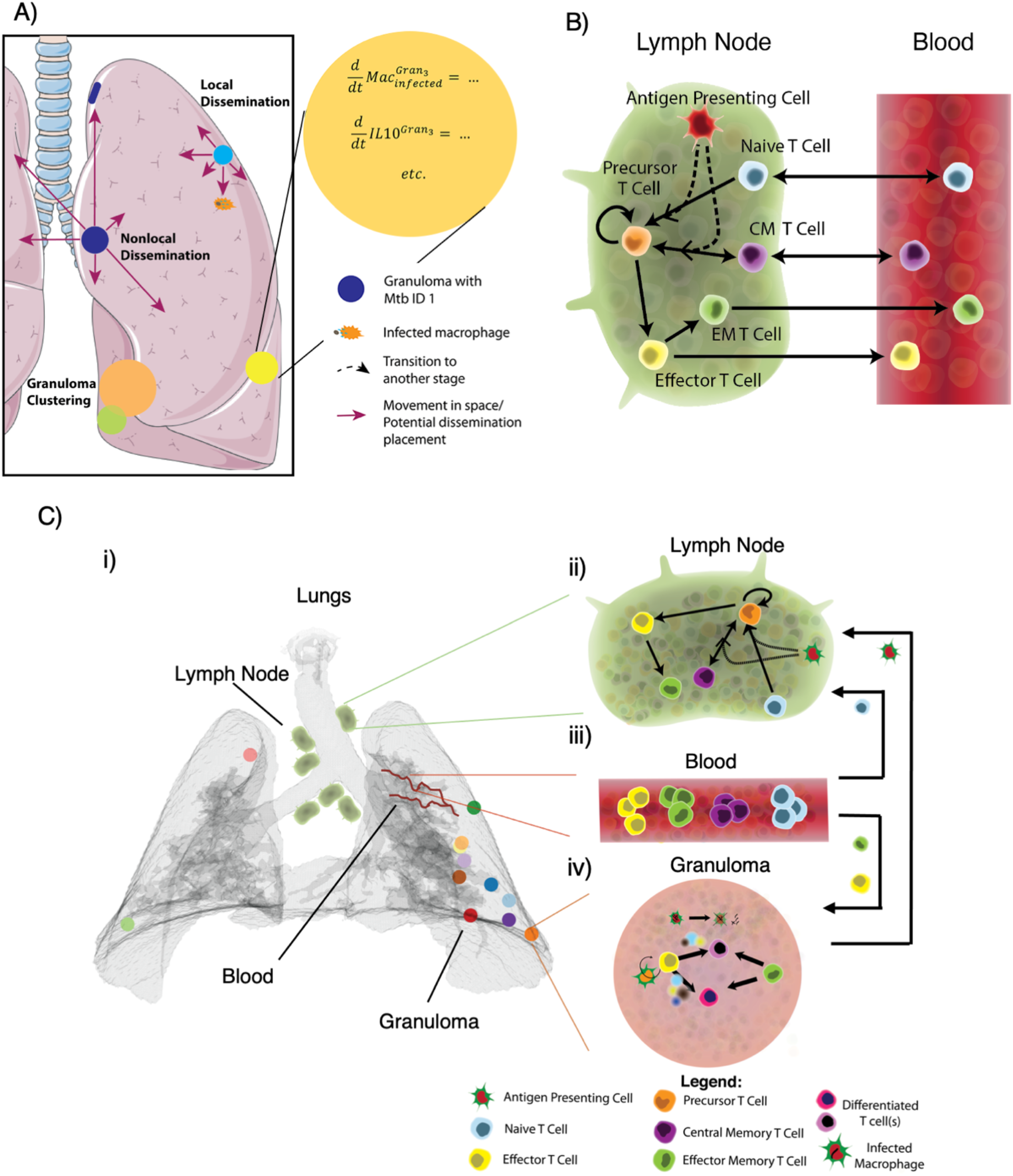
HostSim multi-scale modeling framework. (A) Multiple lung granuloma (*MultiGran*) model conceptual framework. Adapted from Figure 2 in (36). B) The blood and lymph node (LN) model that tracks multiple T cell phenotypes across LN and blood compartments. Adapted from Figure 2 in (38). C) i) *HostSim* model schematic showing lungs (gray), separate granulomas (various colored circles), lung draining lymph nodes (green near trachea), and conceptual lung vasculature (red curves). (ii) Antigen presenting cells travel from lung granulomas to lymph nodes to initiate T cell priming, proliferation, and differentiation. T cells travel from lymph nodes into (iii) blood and re-enter lung granuloma environments (iv) continuously over time to participate in bacterial killing and containment within the granuloma.

Our model is called *HostSim* as we consider a simulation of an entire primate host during Mtb infection; however, our *in silico* “hosts” are comprised solely of lungs, LN and blood. These three physiological compartments comprise the majority of dynamics that occur during pulmonary TB (39,40). Other organs and body system are also involved during extrapulmonary TB, including liver, brain, and other extrapulmonary sites. We believe that focusing this study on pulmonary TB is without loss of generality, and that adding in those other body sites would serve to fine tune our predictions to other clinical outcomes of TB.

Each virtual host includes multiple granulomas with separate parameter values, and a single parameter set for the LN and blood. The assumption that granulomas within the same host have separate parameter values is supported broadly by both modeling and experimental studies that have shown each granuloma within a host evolves independently (9,10,25,26,29,36,41,42).

### Modeling multiple lung granulomas across time - MultiGran

In a recent study, we built a novel hybrid agent-based model that describes the development of multiple lung granulomas known as *MultiGran* (36). In this model, each granuloma acts as an agent, placed stochastically within the boundary of a 3-dimensional lung environment (Figure 1A). To create this ‘virtual lung’ we used a CT scan from an uninfected NHP (36) as the three-dimensional lung architecture upon which multiple granulomas develop across time (translating the *x,y,z* coordinates from a CT scan to our computer model (36)). Simulations begin with inoculation of multiple bacteria into the lung environment. A granuloma is initialized when each Mtb is placed within the lung environment, as NHP studies have shown that each Mtb bacterium can form a unique granuloma (9,10).

Briefly, the development of each individual granuloma “agent” is captured by a system of ODEs that tracks bacterial, macrophage, T cell, and cytokine dynamics. To describe the role of the innate immune response within a granuloma, we track resting, infected and activated macrophages as well intracellular and extracellular bacterial populations. To capture the impact of the adaptive immune system, we track primed CD4+ and CD8+ T cell populations. Primed CD4+ T cells can differentiate into effector Th1 or Th2 populations and primed CD8+ T cell populations can differentiate into cytotoxic or cytokine producing CD8+ T cell populations. Recruitment of T cells from the blood compartment to granulomas is described in greater detail below. We additionally track concentrations of pro- and anti-inflammatory cytokines within each granuloma, including IFN-γ, TNF-α, IL-10, IL-4 and IL-12. *MultiGran* only included the primed and differentiated T cell populations described above; but we now include effector memory T cells to be consistent with experiments that have shown effector memory T cells are present within the granuloma environment (43–45). Thus, we expanded the set of ODEs representing each single granuloma in *MultiGran* (36) to include CD4+ and CD8+ effector memory T cell subpopulations. Briefly, we assume effector memory cells are recruited from the blood to granulomas according to the inflammatory profiles of granulomas (see Linking models section below for further detail). Once at the site of the granuloma, effector memory cells differentiate into T cells that exhibit effector functions (45–47).

Granulomas within *MultiGran* can sterilize bacteria, control bacterial growth over time, or exhibit uncontrolled bacterial growth. Granulomas can also disseminate, spreading bacteria locally or non-locally (Figure 1A). Local dissemination events initialize a new granuloma near the disseminating granuloma whereas non-local dissemination initializes a new granuloma randomly within the lung environment. Model equations and details are in the Supplementary Materials, which includes a list of all parameters, definitions, and ranges.

### Lymph node and blood models

In previous work, we captured LN and blood cellular dynamics following Mtb infection or vaccination using a two-compartment mathematical model (26,38,48). Briefly, we track Mtb-specific and Mtb-non-specific CD4+ and CD8+ naïve, effector, effector memory, and central memory T cell responses using a compartmentalized system of 31 non-linear ODEs (Figure 1B). We represent Mtb-specific T cells as a generic class of antigen-specific cells across time. In the LN, T cells are tracked as counts across time, whereas in the blood, the cells are tracked as a concentration (cells/μL) because experimental data on blood T cells is often presented as a concentration. Supplementary Materials gives the list of all parameters, definitions, and ranges for the blood and LN models.

### Creating the multi-scale model: Linking the lung model (MultiGran) and the lymph node model

T-cell priming, proliferation and differentiation begins in the LN when an antigen-presenting cell (APC) travels from lungs to LN and interacts with a Mtb-specific T cells. In mice, this process does not begin until 9-13 days after inoculation (16,40), but serial positron emission tomography coupled with computed tomography scans (PET-CT) in NHP studies have shown that LNs do not become metabolically active until 2-4 weeks post-infection (39,49,50). Wolf et al. showed that the migration of cells to LN is transient (40), and NHP PET-CT studies revealed that LNs do not increase metabolic activity following 8-12 weeks post-infection during latent infection (49).

We mirror this biological phenomenon in a coarse-grained manner within *HostSim* (Figure 2C). As infection progresses within *HostSim*, we allow infected macrophages within granulomas to act as a proxy for APCs that migrate to the LN beginning ~1-4 weeks post-infection. This assumption is supported by experimental studies and previous modeling has made similar assumptions (36,51,52). We represent the percentage (5-25%) of infected macrophages which will act as APCs and migrate to the lymph node as a parameter that can be varied. This range emerged from calibration, but it is validated by experiments that show only a small fraction of cells can traffic to the LN (51–53). The main migration of immune cells to LNs ceases ~7-14 weeks post-infection, consistent with the NHP PET-CT data (49). However, as TB is a chronic disease, we include stochastic events where a small percentage of cells randomly migrate to the LN every few days. All processes that link lung and LN compartments are events guided by parameters whose initial ranges were estimated from both mouse (16,40) and NHP data (39,49,50). For example, even though we model a single LN compartment, approximately five LNs are involved in NHP and human Mtb infection (50), so we scale all LN T cell counts by a multiple of five when they enter the blood compartment, as done previously (26,37,38).

**Fig 2:**
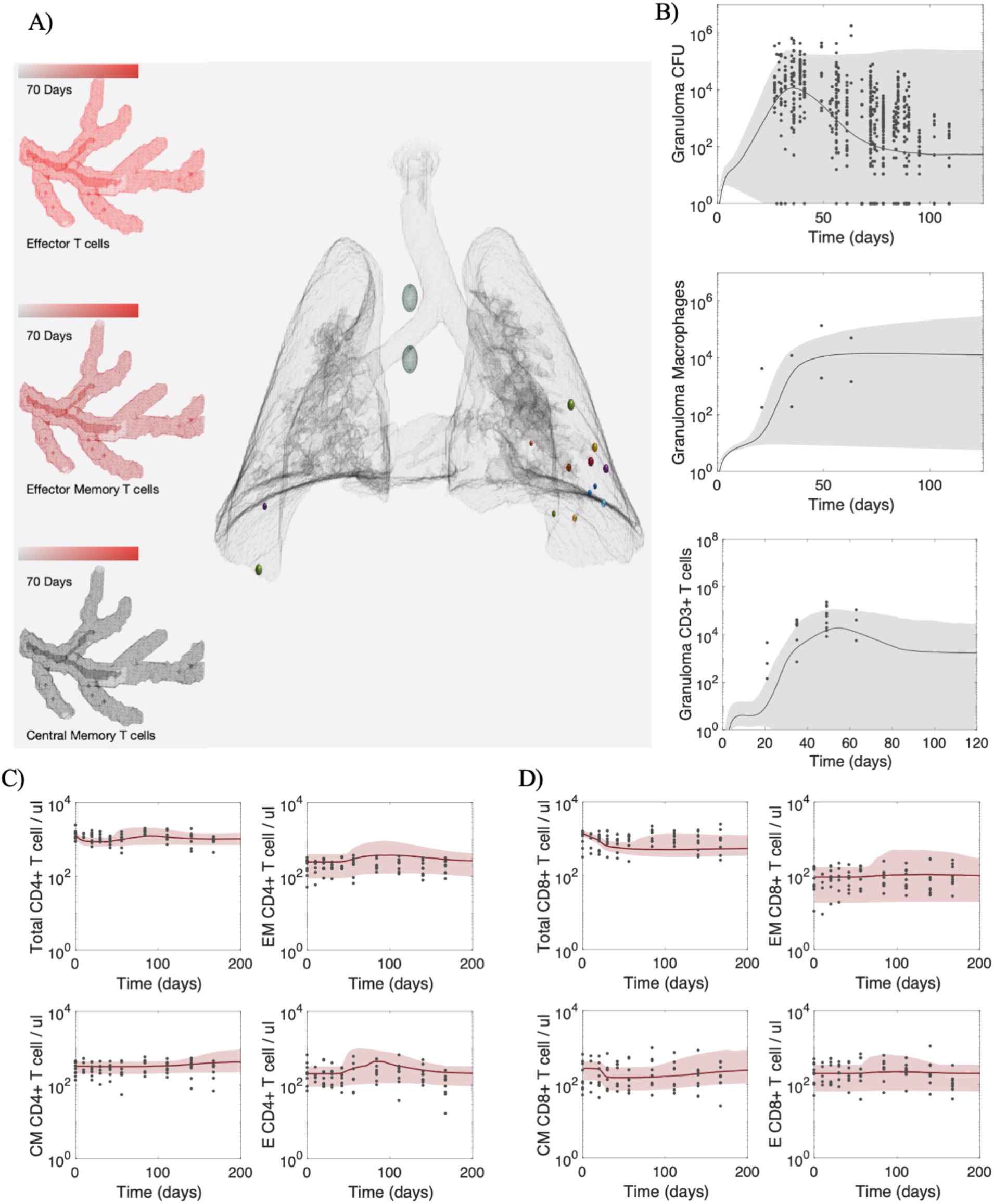
Calibrated HostSim recapitulates dynamics of Mtb infection at both granuloma-scale and host-scale. (A) Snapshot of *HostSim* time-lapse video showing virtual lungs, granulomas, lung draining lymph nodes, and blood cell concentrations for three cell types. Mtb-specific effector, effector memory and central memory T cells numbers within blood are qualitatively captured by a color change across time, from black (very few cells in the blood) to bright red (representing the maximum number of cells of that blood type across the simulation). At day 70, Mtb-specific effector T cells numbers peak, Mtb-specific effector memory T cells are continuing to grow in magnitude, and Mtb-specific central memory T cells have not yet started to differentiate in large numbers. Full time-lapse videos can be found at http://malthus.micro.med.umich.edu/lab/movies/HostSim/. We calibrated *HostSim* to published datasets from NHPs on (B) lung granuloma CFU, macrophage and T cell granuloma numbers from previous studies (26); (C) blood CD4+ T cell data and (D) blood CD8+ T cell data from both simulation and NHP following a single infection event in NHP studies (25,26,64,65). Published NHP study data are shown as black dots across the graphs. For direct comparison, we display simulation data as gray (granuloma outcomes) or red (blood outcomes) clouds that outlines the 1^st^ and 99^th^ percentile across 500 host simulations. Gray and red lines represent the medians of those simulations. Simulations plotted show from day of infection until day 200 post-infection.

### Creating the multi-scale model: Linking the blood model to the lung model (MultiGran)

We also coarse-grain the process of T-cell lung-homing and migration to the sites of granulomas. In *HostSim*, there are three types of blood T cells that are recruited to the granuloma: Mtb-specific effector T cells, Mtb-specific effector memory T cells, and non-specific T-cells. Note, once blood Mtb-specific effector T cells arrive in the granuloma, they are considered primed T cells. Recruitment occurs for both CD4+ and CD8+ T cell lineages.

Each cell type is recruited to each granuloma according to inflammatory signals within our granuloma model. These include counts of activated and infected macrophages, and levels of the pro-inflammatory cytokine TNF, consistent with experimental data and previously presented models (25,37,54–57). We calculate the number of Mtb-specific effector T cells that will be recruited from the blood to the *i^th^* granuloma, *granuloma_i_*, per time step according to the following equation, as outlined in our previous modeling work (33,36,58):

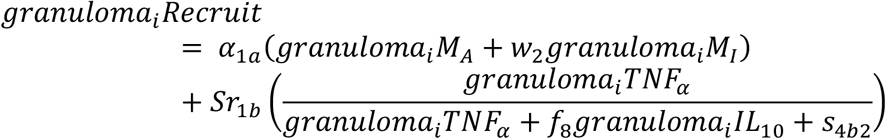

Where *α_1α_, w_2_, Sr_1b_, f_8_, s_4b2_* are granuloma-specific parameters (see Supplementary Material Table 1). Effector Memory T cells are recruited similarly to each granuloma, but recruitment is performed proportional to the level of TNF-α within the granuloma (see Effector Memory T cell granuloma equations in Supplementary Material). We assume different mechanisms of recruitment between these T cell phenotypes arises due to known differences in migration of effector memory and effector T cells to non-lymphoid sites, such as the lung (reviewed in (58)). Altogether, numbers of macrophage and inflammatory cytokine levels act as a proxy within our model for chemotactic and adhesion molecules acting within a granuloma that attract T cells to the site. We perform recruitment for each granuloma at every timestep within the model, i.e. once per 24 hours. At each timestep we update the blood cell numbers by subtracting the summed granuloma recruitment for each cell type, according to the following general form:

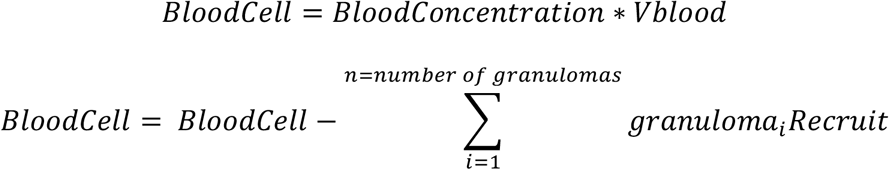

where blood cell concentrations (cells/μL) are converted to blood cell numbers prior to entering the granulomas. Vblood is equal to 3.6×10^5^ μL, a well-established value in the literature that represents the volume of blood (26,37,38,60). This parameter is used to scale cells when they traffic between the blood and the lung or LN compartments. This type of volumetric scaling is standard in compartmental modeling (61).

During very early timesteps following inoculation, granulomas may occasionally attempt to recruit more Mtb-specific T cells than are physically available within the blood compartment. Should this happen, recruitment cell counts are obtained by normalizing the corresponding blood concentrations, such that the magnitude of cell recruitment is proportional to the blood concentration. In general, our assumption that more inflammatory granulomas are able to recruit larger quantities of T cells is consistent with previously presented models and experimental data (25,26,33,42,54).

### Calibrating HostSim to multiple datasets

After construction of *HostSim*, we calibrated the model to estimate model parameter values. An effective strategy to calibrate a complex, multi-scale and multi-compartment system is to calibrate to multiple datasets, thereby reducing the likelihood of parameter overfitting (62). We utilized our previously published protocol for calibrating complex systems to biological data, *CaliPro* (63), to generate a range of calibrated parameter values.

Using *CaliPro*, we simultaneously calibrated to biological datasets across multiple biological scales. We calibrated the single granuloma ODE model to previously published T cell and macrophage datasets from 28 NHP granulomas across 70 days and a bacterial CFU dataset for 623 granulomas from 38 NHPs across 120 days (25,26,64,65). At the whole-host scale, we calibrated the lymph node and blood compartment to a previously published T cell dataset from 26 NHPs across 200 days (26). Each time point within these data sets includes multiple data points, such that the experimental data illustrates a heterogenous range of potential outcomes (Figure 2 B, C & D).

We determined initial parameter ranges for each model parameter based on experimental values from literature, as well as previous single granuloma ODE models, previous lymph node and blood ODE models, and our previous work in modeling multiple granulomas (33,38,58,66). In this modeling framework, some of the parameter values are constrained (such as rates of bacterial killing or cellular death rates) and were not as widely varied as others. We utilized a Latin hypercube sampling (LHS) scheme to sample 500 times within the initial parameter space, thereby creating 500 unique simulations of *HostSim* (i.e. generating 500 unique virtual hosts). We then use *CaliPro* to refine and resample this wide initial parameter space in an iterative manner.

*CaliPro* requires users to explicitly define a *pass set* – this is an automated criterion for which the model simulations can be considered calibrated. We specify a pass set as the simulations that fall within the range bounded by an order of magnitude on either side of the minimum and maximum experimental data point for every time point across each of the experimental outcomes. The experimental data range includes over four orders of magnitude (Figure 2B), therefore our pass set definition was selected since it encapsulates the general behavior of the experimental datasets we are using for calibration and will not remove simulations that are within the same order of magnitude as experimental data points. Additionally, we know that the long-term behavior of bacterial numbers in granulomas are fairly stable without intervention (9), and thus we set an upper bound at 36000 bacteria for days 90-200 as a specific criterion for calibration of this outcome. If the simulation value for bacterial numbers eclipse this bound within those days, the simulation does not belong to the pass set, even if the granuloma T cells and macrophages all lie within the bounds of the experimental data. In an iterative manner, *CaliPro* redefines the parameter ranges for each parameter according to the pass set simulations and reruns the model, comparing against the experimental data until calibration is considered complete (a pre-defined user input). For *HostSim*, calibration was considered complete when 90% of simulations belonged to the pass set. Supplementary Material Table 1 lists the calibrated parameter ranges for each varied parameter.

### Sampling parameter space to create HostSim virtual hosts

We sample from our calibrated parameter space to create each unique *HostSim* virtual host. Each host is composed of one parameter set that guides the LN and blood ODE model and one unique parameter set for each granuloma within the virtual host. When we generate a population of virtual hosts, we sample uniformly from our parameter space for the LN and blood model according to an LHS scheme (67) and select a single parameter set for each host. When sampling the granuloma ODE parameter space, we again utilize an LHS scheme to select an initial point in parameter space for each host, and then sample each granuloma parameter set for that host from a normal distribution. For each parameter, the mean of the normal distribution is set as initial point in parameter space as selected by LHS and σ is set to be equal to one-quarter of the parameter’s calibrated range. We sample the granuloma parameter space once for every granuloma that is initialized within an individual at the time of inoculation (this number varies depending on the inoculation dose used in the virtual experiment). Together, the granuloma parameter set and the LN and blood parameter set are the inputs for a single virtual host simulation.

### Using bacterial numbers as a proxy for clinical classifications in HostSim

To explore the range of possible host-scale outcomes in *HostSim*, we sample from our calibrated parameter space and generate a virtual population of 500 unique hosts. Each individual simulation begins with an inoculation dose of 10 CFU, stochastically placed within the lower left lung lobe to seed the formation of 10 unique granulomas. We start each simulation with 10 CFU to be consistent with the inoculation of NHPs, which inoculate ~10 CFU to begin experiments (68).

Each virtual host in the population is simulated for 200 days. At 200 days, we delineate clinical classifications across the population of 500 virtual hosts according to the total lung CFU per host. We calculate the total lung CFU by summing the individual granuloma CFU for all granulomas within a host at each time point. We use the following cutoffs for clinical classification: TB eliminators: total lung CFU<1; Active TB cases: total lung CFU > 10^5^; LTBI: all other virtual hosts. We establish the threshold between active TB cases and LTBI cases in *HostSim* to be consistent with NHP studies that show that total bacterial burden in active TB cases is significantly higher than that of LTBI monkeys, although the same study did show a small number of active cases with a bacterial burden similar to that of latent NHPs (see Discussion and (21) for more detail). Finally, we select 200 days (~7 months) post-infection for clinical classification in order to be consistent with NHP studies that classify animals 6-8 months following infection (69).

In the dose inoculation studies, we use the same virtual population of 500 hosts, but run 25 separate virtual experiments and vary the inoculation dose from 1-25 CFU. Thus, depending on the study, hosts begin the simulation with 1 to 25 unique granulomas. At the conclusion of the simulation – day 200 – we use the same thresholds of total lung CFU for determining clinical classifications across all hosts.

### Uncertainty and Sensitivity Analysis

We quantify the importance of host-scale and granuloma-scale mechanisms involved in infection outcomes using statistical techniques known as uncertainty and sensitivity analysis. As mentioned above, we efficiently sample our multi-dimensional calibrated parameter space using LHS algorithms to generate 500 individual virtual hosts. We then determine correlations between model outputs and parameter values by using Partial Rank Correlation Coefficient (PRCC), a common method for determining correlation-based sensitivity (67).

Sensitivity analyses of multiscale models can be difficult (70). ‘All-in-one’ sensitivity analyses are one method for exploring relationships between model parameters and outcomes by treating the full model as a black box and varying all parameters. In particular, ‘all-in-one’ sensitivity analyses are not always sufficient for understanding relationships between model parameters and outcomes, especially when a model is sufficiently complex and composed of multiple compartments or sub-models, as is the case with *HostSim*. As reviewed in (71), an ‘all-in-one’ sensitivity analysis can be paired with an intra-compartmental model approach to provide comprehensive understanding of the model behavior across scales.

We present results from two separate sensitivity analyses. First, we vary parameters across the whole-host scale and granuloma-scale physiological compartments to create 500 unique virtual hosts. Each virtual host in this population includes multiple granulomas with separate parameter values. We perform an ‘all-in-one’ sensitivity analysis across these 500 virtual hosts to identify significant associations between parameters and whole-host clinical outcomes in TB (i.e., LTBI or active TB cases).

Next, to perform an intra-compartmental analysis, we select two representative hosts – one host that was classified as an active TB host and one that was classified as a TB eliminator according to their total lung CFU at day 200. For each representative host, we rerun the simulation 500 times, varying only granuloma-scale parameters while fixing the blood and LN parameters (Supplementary Material Figure 1 displays granuloma CFU trajectories of each set of 500 simulations). From each set of simulations, we calculate PRCC values to identify associations between granuloma-scale parameters and granuloma CFU at day 200. We performed False Discovery Rate test corrections (72) on all reported significant parameters.

### Pro- and anti-inflammatory profiles of HostSim granulomas

We present a unitless measure that represents the ratio of pro- and anti-inflammatory cytokines for granulomas within *HostSim*. Cytokine units in *HostSim* granulomas are picograms per microliter, a measure that is consistent with previously published models of cytokine levels in granulomas (33,58,73). However, to investigate relative ratios of pro- and anti-inflammatory cytokines within each *HostSim* granuloma, we calculated the common logarithm (logarithm with base 10) of the IL-10, TNF-α and IFN-γ cytokines and plotted these values in a 3-dimensional scatterplot. This allows for a comparison of granuloma inflammatory profiles, across orders of magnitudes of cytokine concentrations within the granuloma environment.

### Model analysis tools and simulation environment

Model code and preliminary data analyses are written in MATLAB (R2020a). We solve the systems of ODEs using MATLAB’s ode15s stiff solver, using a timestep of one day. At the end of each timestep, we perform cell recruitment and update granuloma cell, cytokine, and bacterial states as well as lymph node and blood cell concentrations. A single *in silico* individual simulation across 200 days of infection time can be performed on a 2-core laptop in approximately 5 minutes. We wrote bash scripts to submit multiple runs of *HostSim* on compute clusters. We perform post-processing statistical analysis, graphing and movie rendering within MATLAB (R2020a).

## Results

### HostSim recapitulates in vivo granuloma-scale and host-scale dynamics

We calibrate *HostSim* to published datasets from NHPs across multiple scales following a single primary infection event. We utilized *CaliPro*, our protocol to define and perform calibration for computational models (63). *CaliPro* identifies a parameter space where each varied parameter has a range of values that correspond to a range of outcomes that match experimental datasets. For this work, the experimental data come from published NHP studies (10,25,36,65). Our *HostSim* website shows calibration datasets and references for each dataset (http://malthus.micro.med.umich.edu/lab/movies/HostSim/).

When sampling parameter sets within our calibrated parameter ranges, *HostSim* matches both the range of experimental outcomes and the dynamics outlined by datasets of primary Mtb infection derived from published NHP studies (Figure 2). At the granuloma scale, *in silico* granulomas from *HostSim* simulations are able to replicate NHP granuloma CFU, T cell and macrophage dynamics across time (Figure 2B, experimental data from previously published NHP studies (10,25,36,65)). Granuloma CFU peaks at approximately 35 days as macrophage and T cell counts increase. Following the peak, CFU, macrophage and T-cell counts correspondingly stabilize across time. At the host scale, *in silico* blood cell counts replicate NHP blood CD4+ and CD8+ T cell data across time (26). Following infection, there is a slight peak in overall effector and effector memory T-cell types that precedes a growing number of central memory CD4+ and CD8+ T cells. (Figure 2 C&D). Across multiple scales, *HostSim* presents a ‘virtual host’ model of the immune response to Mtb infection.

### Emergent HostSim behavior across a virtual population matches spectrum of tuberculosis

Humans present a spectrum of clinical outcomes in TB, including (but not limited to) complete elimination of infection, latent infection, and active TB disease (3). Work in NHPs have shown that total bacterial burden is associated with clinical outcomes. Specifically, total bacterial burden in active TB cases is significantly higher than that of LTBI monkeys (21). *HostSim* exhibits similarly heterogenous host-scale outcomes (Figure 3).

**Fig 3:**
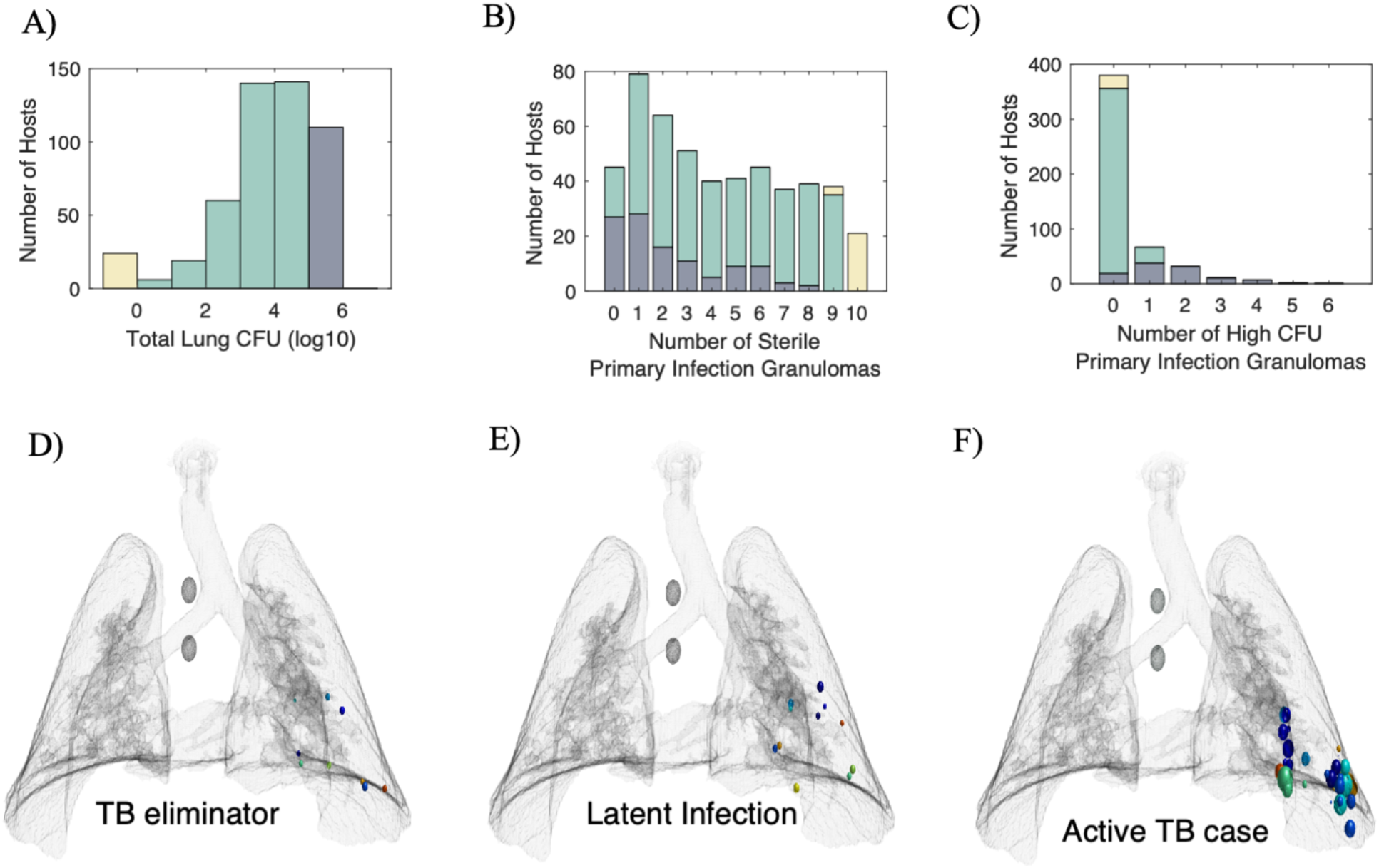
HostSim exhibits a spectrum of whole-host outcomes across a population of 500 virtual hosts. (A) Histogram displaying the total lung CFU per host at day 200 across our virtual population of 500 hosts. We delineate the virtual population into three groups: TB eliminator (yellow), LTBI (green), or active TB cases (dark blue) according to the total Lung CFU. (B) Stacked bar chart displaying the number of sterile granulomas per host across TB eliminator, LTBI, or active TB cases. (C) Stacked bar chart displaying the number of high CFU granulomas per host across clinical scale outcomes: TB eliminator, LTBI, or active TB cases. (D, E, F) *HostSim* snapshots display virtual lung architecture and granuloma locations for representative TB eliminator, LTBI and active TB cases at day 200 post-infection.

To explore ranges of host-scale outcomes in a virtual host study, we sample from our calibrated parameter space to generate a virtual population of 500 unique hosts. Each simulation begins with an inoculation dose of 10 CFU (selected to be consistent with inoculation of NHPs (68)), thereby starting the formation of 10 individual granulomas within the lung environment. Simulations run for 200 days. We calculate the total lung CFU by summing the individual granuloma CFU for all granulomas within a host.

Across our virtual population of 500 virtual hosts, the total lung CFU per host spans several orders of magnitude, from 0 CFU (infection elimination) to 10^6^ CFU (Figure 3A). We delineate our virtual population into 3 groups according to their total lung CFU at day 200, analogous to the clinical classifications of NHPs 6-8 months following primary infection (69). We use the following cutoffs for classification: TB eliminators: total lung CFU<1; Active TB cases: total lung CFU > 10^5^; LTBI: all other virtual hosts. Across our 500 virtual hosts, there are 24 TB eliminators, 110 active TB cases, and 366 LTBI individuals. Snapshots from representative simulations of these diverse outcomes are displayed in Figure 3D, 3E & 3F.

After classifying the virtual hosts by total lung CFU, we looked at two additional statistics. First, we examined the number of sterilized granulomas across the three different clinical classifications (Figure 3B). Our model predicts that ~75% of active TB cases include at least one sterile granuloma. This finding is validated by a previously published NHP dataset, which showed 11 out of 13 monkeys with active TB had at least a single sterilized granuloma (9).

Second, we looked at the number of virtual hosts that have individual granulomas with a high bacterial burden (defined as granulomas with 5×10^4^ CFU or higher; Figure 3C). As expected, all TB eliminators and the majority of LTBI virtual hosts do not contain a granuloma with a high bacterial burden. However, we see approximately 8% of our LTBI classified hosts include one high CFU granuloma. These cases indicate that our model may have the potential to capture incident or subclinical TB and may explain the spectrum nature of TB disease as these individuals could be more likely to reactivate or progress to active disease (5).

### A multi-scale sensitivity analysis reveals adaptive immunity drives clinical classification and innate immunity impacts granuloma-scale outcomes

We next use the model to investigate mechanisms that drive host-scale clinical outcomes. Using uncertainty and sensitivity analysis, we can identify these driving mechanisms across multiple scales of interest. First, we perform an ‘all-in-one’ sensitivity analysis (see Methods) on clinical classifications (see Figure 3) across the 500 virtual hosts from our calibrated parameter space. Table 1 highlights parameters found to be significantly correlated (p<0.05) with each clinical classification from our PRCC analysis. Not surprisingly, we find that elements of adaptive immune responses within LNs are main drivers of whole-host clinical outcomes. Specifically, the differentiation and proliferation of T cells within LNs are significantly associated with clinical scale outcomes (i.e. active TB, LTBI or TB eliminator). The significant, positive association between T-cell proliferation in LN and clinical classification at the whole-host scale represents an inter-physiologic compartmental effect – not only do LN parameters influence T-cell counts within the LN, but they influence whole-host scale clinical outcomes as well. Further, both Mtb-specific CD4+ and Mtb-specific CD8+ T cell parameters in the LN impact clinical-scale outcomes, lending further support to emerging studies showing the importance of CD8+ T cells in TB (45,68,74).

**Table 1:** Parameters identified as significant from sensitivity analysis. For each analysis, parameters shown here have a PRCC absolute value of ρ > 0.1 and p-value<0.05. Parameters listed as associated with clinical outcomes are the result of our ‘all-in-one’ sensitivity analysis. Clinical-scale classifications were assigned a value of 0 (active TB case), 1 (LTBI) or 2 (TB eliminator) to calculate the PRCC value for each parameter. Parameters listed as associated with granuloma CFU were the result of our intra-compartment analysis. These parameters were significantly correlated with granuloma CFU at day 200. PRCC values are listed in Supplementary Material.

| Parameters associated with clinical-scale outcomes | Description of parameters<br>‘All-in-One’ sensitivity analysis |
| --- | --- |
| LN_k13 | Precursor CD8+ T cell proliferation within the lymph node |
| LN_k14 | CD8+ T cell differentiation to CD8+ effector T cell in lymph node |
| LN_k4 | Precursor CD4+ T cell proliferation within the lymph node |
| LN_k5 | CD4+ T cell differentiation to CD4+ effector T cell in lymph node |
| Parameters associated with granuloma-scale CFU outcomes | Description of parameters<br>Intra-compartment sensitivity analysis |
| k2 | Resting macrophage infection rate |
| c9 | Likelihood of resting macrophages to phagocytize bacteria |
| N | Carrying capacity of intracellular bacteria within macrophages |
| k17 | Max rate of infected macrophage death from intracellular bacteria |
| k18 | Extracellular bacterial killing by resting macrophages |
| k14a | Fas:FasL induced apoptosis of MI |
| alpha11 | IL-4 production from primed T cells |

To explore the drivers of granuloma-scale variation within a host, we perform an intra-compartmental sensitivity analyses (see Methods) focusing solely on which granuloma-scale parameters are associated with granuloma CFU at day 200. This allows us to identify how granuloma scale parameters may contribute to heterogenous granuloma CFU outcomes within a host when blood and LN parameters are held fixed (PRCC values are given in Supplementary Material). The bottom half of Table 1 lists mechanisms that we identified from both the adaptive and innate immune responses. Multiple parameters that dictate macrophage behavior were identified as key drivers of granuloma CFU. Additionally, adaptive immune response parameters were also associated with reduced granuloma CFU (i.e., Fas:FasL cell death in Table 1).

Altogether, the results from our ‘all-in-one’ sensitivity analysis as well as our intra-compartmental analyses predict that while the adaptive immune response in LNs drive clinical-scale outcomes, the innate immune system does play an important role within a host by contributing to heterogeneity of granuloma CFU, as observed within humans and NHPs.

### Infection outcomes of virtual hosts are dose dependent

In humans, a relationship between inoculation dose and severity of clinical disease has been hypothesized (75–77). To explore this in our virtual hosts, we performed a set of inoculation dose experiments using *HostSim*. We reran our virtual population of 500 hosts through 25 simulated experiments. For each experiment we re-simulated the 500 virtual hosts with identical random seeds and parameter sets, varying only dose inoculum from 1 to 25 CFU. Figure 4 displays the total lung CFU and clinical classification of those 500 hosts at day 200 following each of the 25 experiments.

**Figure 4:**
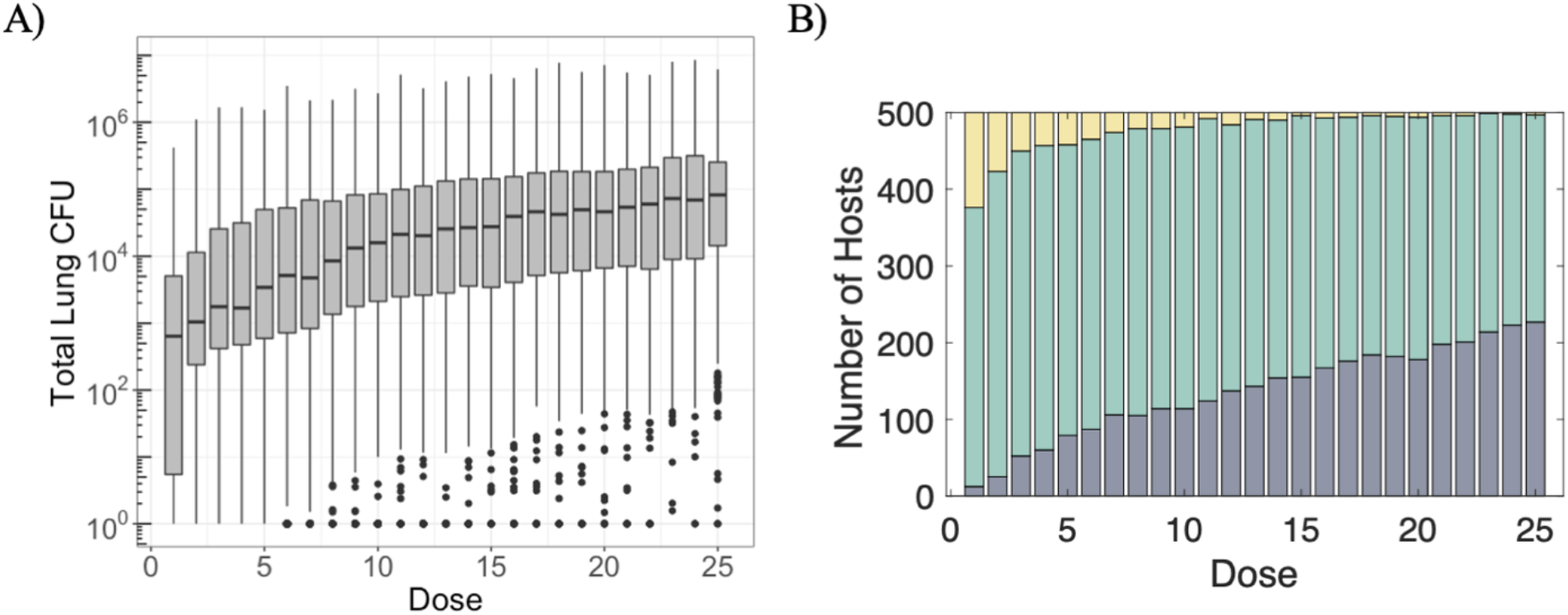
Infection outcomes at day 200 post-infection across a population of 500 virtual hosts are dose dependent. A) Distribution of total lung CFU per host among the virtual population for the 25 inoculation dose experiments. Total lung CFU is calculated by summing CFU across all granulomas in a single host. B) Stacked bar charts display virtual host clinical-scale outcomes based on total lung CFU per host for the 25 inoculation dose experiments. Bar chart colors are the same as Figure 3 - TB eliminators (yellow), active TB cases (dark blue) or LTBI (green).

As dose inoculum increases, the median lung CFU for the population of 500 hosts (at day 200 post-infection) increases; however, the model predicts a range of outcomes across the population for each inoculation dose (Figure 4A). For example, among the 500 hosts inoculated with 25 CFU, a few hosts had low levels of CFU within the lung (CFU <100). Conversely, after a dose inoculum of 1 CFU, some hosts still exhibited considerable infection, with total lung CFU > 10^5^.

For each of the 25 dose experiments, clinical classifications of the virtual hosts based on the total lung CFU at 200 days post-infection are shown in Figure 4B. We delineated the virtual population into three clinical outcome groups as above, where TB eliminators have a total lung CFU<1, active TB cases have a total lung CFU > 10^5^ and all other hosts are classified as LTBI. After an inoculum of 25 CFU, ~55% of the simulations are classified as LTBI and ~45% are classified active TB cases at day 200 (Figure 4B). Thus, *HostSim* predictions agree with human association studies (75–77) suggesting TB disease severity is dose dependent.

### The fates of individual granulomas are heterogeneous within hosts

In both human and NHP studies, individual granulomas within a single host can present a heterogeneous array of morphological, pathological, and immunological outcomes (41,78–80). In NHP studies, even granulomas within active TB monkeys can exhibit sterilization (9,21,81). Similarly, within individual hosts across our virtual population of 500 hosts, we identify a range of granuloma-scale outcomes, from total sterilization to uncontrolled bacterial growth. Figure 5 displays individual granuloma CFU trajectories from five representative hosts ranging across different clinical-scale outcomes: TB eliminator, LTBI and active TB, respectively. Within-host variation is apparent in all hosts, but we highlight that host #5 has both sterilized and disseminating granulomas present. Dissemination occurs when bacteria escape one granuloma and seed the formation of another granuloma elsewhere in the lung environment (36). Dissemination granulomas can be identified when a new CFU trajectory begins at any timepoint after the initial infection (c.f. Figure 5B host #5). However, dissemination does not only occur in active TB hosts; we also note a dissemination event occurred in host #3 (Figure 5B), a virtual host that is still classified as LTBI according to our established criteria outlined in Figure 3A and Methods.

**Fig 5:**
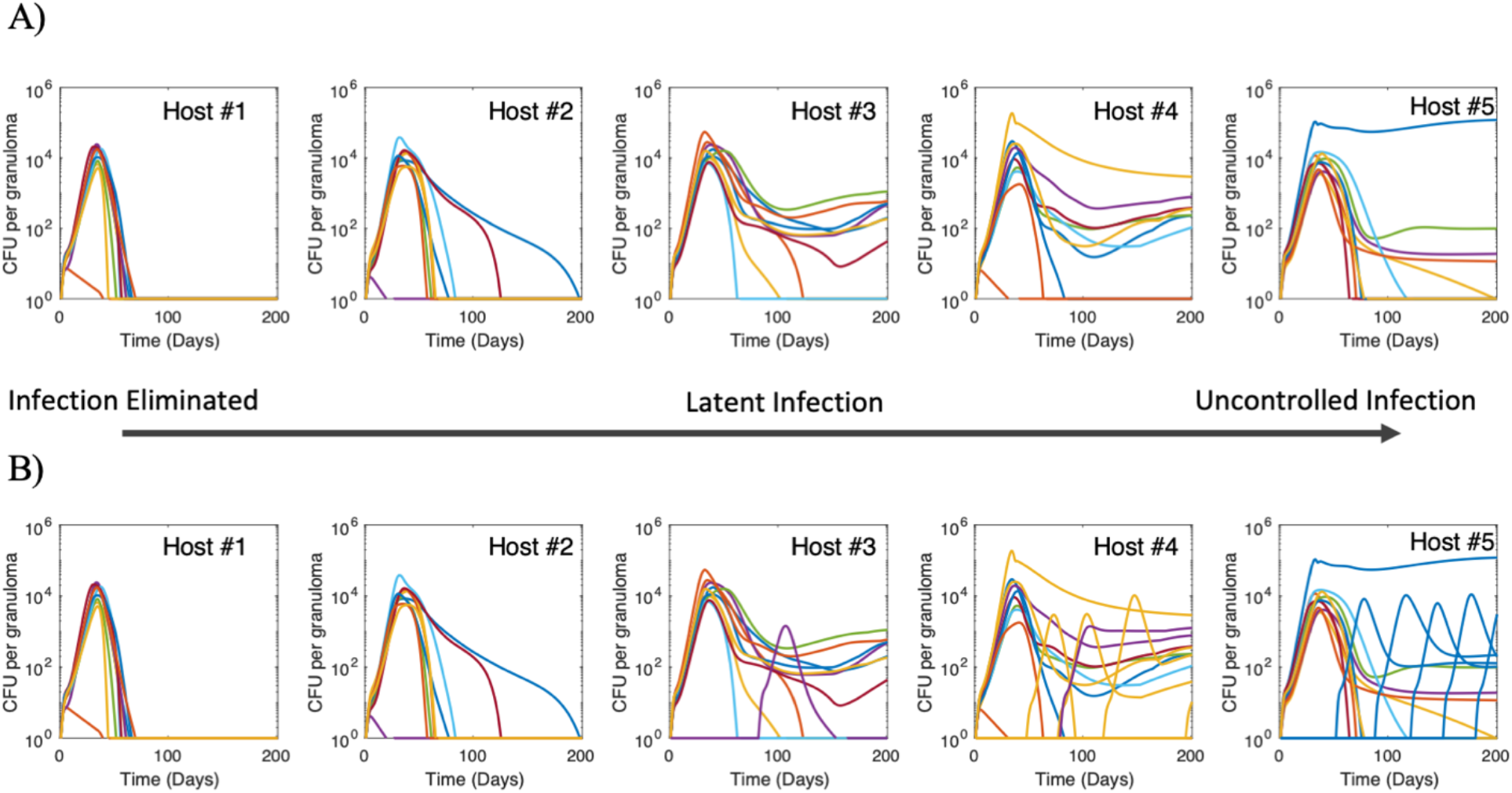
*HostSim* exhibits spectrum of granuloma-scale outcomes within hosts. 500 virtual hosts were simulated to create our population, as shown in Figure 3. We identified 5 representative hosts that exhibited a spectrum of whole-host outcomes (elimination, control and uncontrolled infection outcomes). Each graph is an individual host – the same five hosts are shown in (A) and (B). Each curve represents the CFU in a single granuloma within the host over time. Sterilization of an individual granuloma can be seen when CFU reaches 0 at any time post-infection. Dissemination occurs when a new curve begins at any time after initial infection. Dissemination granuloma CFU trajectories are colored to match the granuloma from which they disseminated. (A) Individual granuloma CFU trajectories for primary infection granulomas only within the 5 representative virtual hosts. B) Primary infection and dissemination granuloma CFU trajectories across the same 5 virtual hosts. Note that in the far-right of panel B), one granuloma (blue CFU trajectory) incurred multiple dissemination events, spurring the formation of multiple new granulomas across time.

For the majority of hosts across our virtual population, the fates of primary infection granulomas are sufficient to delineate clinical-scale outcomes at day 200. Out of the 500 *in silico* hosts, only 8 hosts (~2%) are reclassified as active TB cases when considering both primary infection granuloma and disseminating granuloma bacterial burdens. That is, the outcomes of dissemination granulomas are often not necessary to classify clinical cohorts within *HostSim*. This prediction suggests that the fate of host clinical-scale outcomes is determined at early stages of infection, even prior to dissemination events that occur after inoculation.

### Early events across multiple scales during infection are predictive of TB clinical outcome

Early events in Mtb infection are thought to impact late-stage clinical-scale outcomes (13,15,69,82). However, this is a difficult relationship to investigate clinically or experimentally. Once an animal is necropsied there is no way to know *a priori* if that animal would have progressed to active or latent infection. *HostSim* provides a tool through which we can relate early events within the lungs and LNs to clinical-scale outcome (TB eliminators, LTBI, or active TB) determined months later across our virtual population of hosts. In the last section we predict that mechanisms operating within granulomas at early stages across multiple scales impact clinical-scale classifications. At the host scale, we investigate relationships between blood and lung immune cell counts. Additionally, we stratify lung T-cell counts by clinical-scale outcomes. At the granuloma scale, we examine the ratio of pro- and anti-inflammatory cytokines within the granuloma.

First, we test whether there is a relationship between levels of immune cells in the blood and within the lung. Figure 6A shows an association between lung and blood levels of T cells at day 50 for four separate T cell phenotypes (Mtb-specific CD4+ effector, effector memory and Mtb-specific CD8+ effector, effector memory) across the 500 virtual hosts. Day 50 was selected as it is typically the height of effector-expansion within in the model, timing that is supported by the NHP granuloma and blood T cell datasets (c.f. Figure 2). Each datapoint is colored according to the simulations’ clinical outcomes at day 200. Note that there is a relationship between numbers of lung and blood CD4+ effector T cells and CD8+ effector T cells (r = 0.5, p < 0.01 and r = 0.61, p < 0.01, respectively). However, by day 200 (Figure 6B), the time point we use for clinical classification, this relationship between blood and lung numbers is less clear (r = 0.3, p < 0.01 and r = 0.14, p < 0.01; for CD4+ and CD8+ effector T cells, respectively).

**Fig 6:**
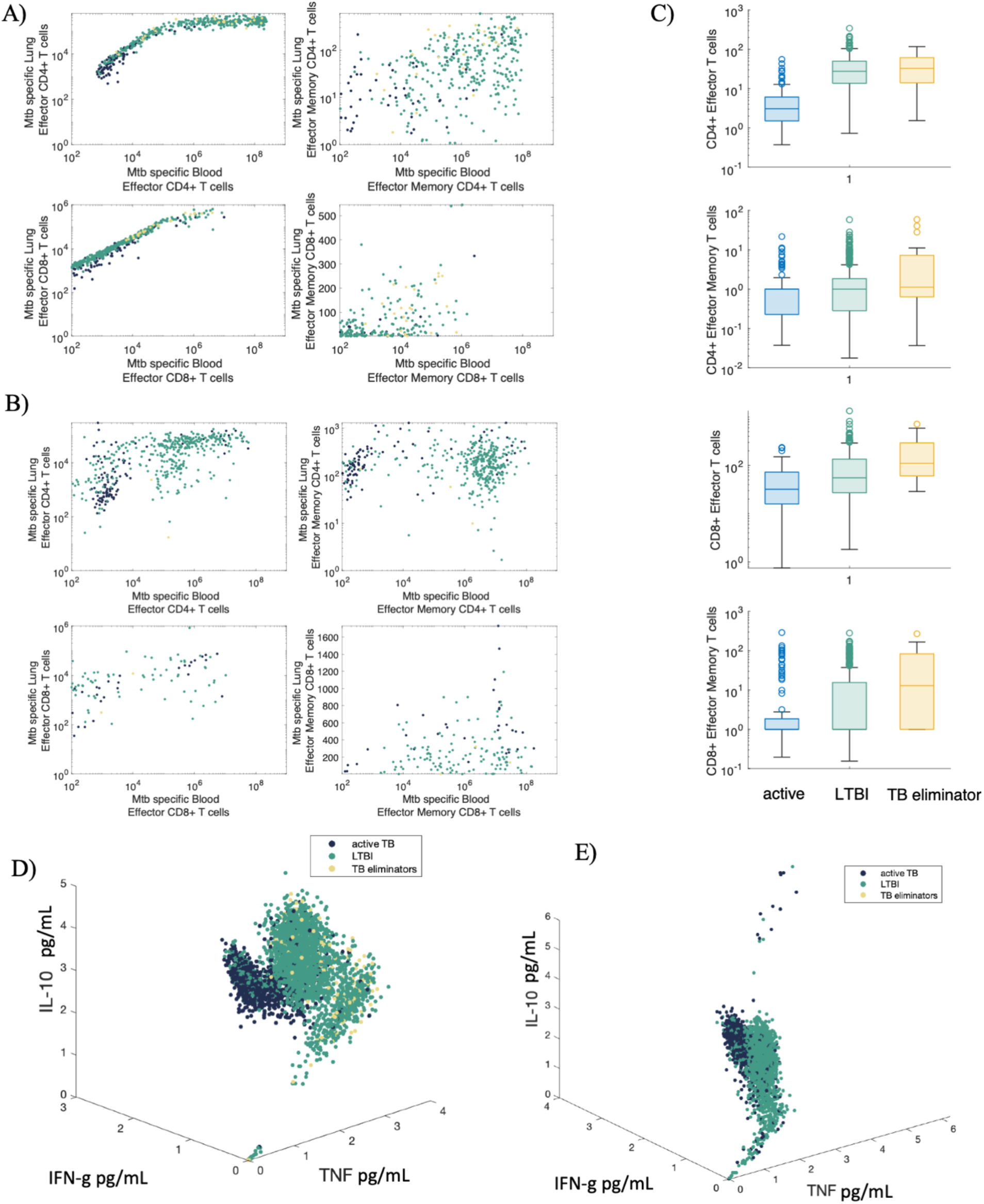
Early events at granuloma-scale and host-scale can predict clinical classifications across a population of 500 virtual hosts. Scatterplots display blood (x-axis) and lung (y-axis) cell counts for Mtb-specific effector and effector memory CD4+ and CD8+ T cells at day 50 (A) and day 200 (B). C) The fold change in numbers of lung T-cells between day 30 and day 40, grouped by clinical classifications at day 200. Each graph displays the fold change for a separate T cell phenotype in the lung. All granulomas from 500 virtual hosts plotted according to relative concentration TNF, IFN-γ and IL-10 cytokine concentrations (pg/mL) on a log scale (see Methods) at day 60 (D) and day 200 (E) colored according to the classification of the host within which the granuloma resides. Across all plots, dark blue = active TB cases, green = LTBI, yellow = TB eliminators.

Second, we identify a fold-change difference in numbers of lung T cells between days 30 and 40 post-infection as indicative of clinical classification 160 simulation days later (Figure 6C). Across the four Mtb-specific T cell phenotypes that are recruited into the lung (Mtb-specific CD4+ effector, effector memory and Mtb-specific CD8+ effector, effector memory), virtual hosts that are classified as TB eliminators typically had a larger fold-change difference between days 30 and 40 than did virtual hosts that are classified as active TB or LTBI cases at day 200 (Supplementary Material Table 2 shows Vargha and Delaney’s A measure for effect size comparisons across all clinical outcomes). Specifically, the median fold change between days 30 and 40 of numbers of Mtb-specific CD8+ effector memory T cells in TB eliminator virtual hosts is approximately 10x larger than that of active TB virtual hosts. We observe a similar difference between LTBI and active TB virtual hosts for numbers of Mtb-specific CD4+ effector T cells. These results suggest that numbers of these cell types have a crucial and early role that impacts clinical classifications made over 150 days later.

Finally, the cytokine profile of granulomas at early time points is indicative of downstream clinical-scale outcomes. Figure 5D shows a three-dimensional scatterplot of pro- and anti-inflammatory cytokine concentrations (pg/mL of IFN-γ, TNF-α, and IL-10) of every granuloma at day 60 across the 500 virtual hosts. Each granuloma data point is colored according to the classification of the host within which the granuloma resides. Note that a cluster emerges wherein granulomas with high levels of IFN-γ, low levels of TNF-α, and low levels of IL-10 are indicative of granulomas that are destined to be within active hosts. By day 200 (Figure 6E), this cluster cannot be as easily separated from the other simulations. Taken together, these predictions suggest that the dynamic balance of pro- and anti-inflammatory cytokines across time (83) could obscure this finding for granulomas sampled at later timepoints.

## Discussion

Tuberculosis is a complex and heterogenous disease. At the host-scale, the disease can manifest across a spectrum of clinical-scale outcomes, including but not limited to TB eliminators, LTBI or active TB (3). Within a single host, individual granulomas are diverse in terms of morphology, immunology and bacterial burden. One of the most highly studied aspects of TB pathology is the granuloma, but a link between granuloma-scale outcomes and whole-host outcomes has yet to be elaborated. Even active TB cases can contain a non-uniform collection of granulomas, wherein a subset of granulomas sterilize bacteria despite a collective failure by the host to rid the body of disease (9). Using experimental studies alone, it can be challenging to identify mechanisms responsible for such heterogeneous outcomes within and across hosts in TB. Mathematical and computational modeling approaches provide powerful tools to link events operating within multiple physiological compartments to host-scale clinical outcomes.

In pursuit of a better understanding of events occurring across multiple-biological scales leading to distinct clinical-scale outcomes, we develop a first-of-its-kind, multi-scale and multi-compartment model of whole-host Mtb infection called *HostSim*. This generalized model is an initial step toward the realization of personalized digital twins in TB research (84,85). We calibrate and validate *HostSim* against previously published, distinct NHP datasets that span cellular, bacterial, granuloma and whole-host scales and make predictions about events that may cause heterogeneous outcomes across multiple scales.

An effective weapon in the global public health battle against TB is identification of robust biomarkers for disease diagnosis and treatment. In TB, there have been many studies and debates regarding both the identification and usefulness of biomarkers (86–93). One barrier to identifying robust biomarkers is the variability in disease outcomes between, and within, hosts at a population scale. In this work, we presented evidence for another relatively unconsidered barrier: biomarkers are transient over time by their very nature. Here we have predicted that the relationship between numbers of blood immune cells and numbers of cells within the lung may only be well-defined at early time points post-infection. Months, or years later, when an individual might present in clinic (94), blood immune cell levels may not accurately reflect events within the lung and therefore may not be a useful compartment to sample when delineating disease status or progression. This reflects a key *HostSim* prediction: recent efforts to identify events in the blood that may correlate with events in lung (23,24) may not be generalizable to every time point for every patient. This prediction is consistent with a recent NHP study that shows blood T-cell responses do not consistently reflect T-cell responses within granulomas (25). These findings are more broadly supported by the idea of a dynamically balanced immune response that occurs across time during chronic infections (83).

In TB animal studies, experimentalists are often unable to know *a priori* if animals necropsied at early time points were destined to be classified as active or latent (69). Using our virtual population of 500 hosts, we are able to show that early events at both granuloma- and host-scales can be predictive of clinical-scale outcomes ~150 days later (at 200 days p.i.). These predictions are potentially useful for experimentalists, who can use analogous experimental techniques (such as serial intravascular staining (95), or IHC cytokine staining of granulomas (96)) to make educated predictions about downstream clinical-scale outcomes. Further, these *HostSim* predictions contribute to a growing body of evidence that suggests early immune events matter in TB (15,82,97).

As the primary intracellular niche for Mtb during both early and chronic stages of infection, macrophages play a central role in TB pathology (98). Recent experimental work has identified Bacille Calmette Guérin (BCG), the only licensed TB vaccine, as a potentially potent innate immune response stimulator by educating macrophage progenitors (99,100). In this work, we used sensitivity analysis techniques to show that parameters governing interactions between Mtb and macrophages at the granuloma-scale are important contributors to the heterogenous granuloma outcomes. Together, these studies and our predictions suggest that macrophages could be viable targets for future therapeutic interventions in TB. This follows as macrophages are crucial cells that sit at the intersection of adaptive and innate immune responses against Mtb.

There are a few limitations to our study and model. First, while we call *HostSim* a whole-host model of Mtb infection, we only represent three physiologically unique compartments (lung, lung-draining lymph nodes and blood). Some of the most progressive forms of TB include extrapulmonary disease (101). As it is beyond the scope of this work, we do not capture the dynamics of extrapulmonary disease with this model, though future work could focus on the dissemination of bacteria into the LN as an initial step to model extrapulmonary disease. Second, while *HostSim* has been developed based on previous modeling efforts and extensive NHP datasets, it does not include all the various cell types present within the granuloma environment (i.e. neutrophils (102,103) or fibroblasts (104)). These cells were not included here primarily because datasets were not as readily available or mechanistic functions of these cells within granulomas are not as well characterized. The *HostSim* modeling framework is flexible and can include these cell types in the future as more data become available about their role in TB granulomas. This limitation extends to the LN and blood compartment models as well, where we do not capture the events of every cell type involved in Mtb infection (i.e., B cells in the LN). Finally, *HostSim* does not capture physical symptoms of TB disease such as coughing or weight loss. Accordingly, we assumed total lung bacterial burden can be used as a proxy for clinical-scale classifications of TB. This assumption is not without precedent. Antibiotic studies in TB frequently use sputum-based assays as a proxy for drug efficacy and assessment of treatment progression in humans (105). Further, NHP studies have shown that total bacterial burden in active TB cases is significantly higher than that of LTBI monkeys, although the same study did show a small number of active cases with a bacterial burden similar to that of latent NHPs (21). Thus, our cut-off for active TB cases (total lung CFU>10^5^) in *HostSim* virtual hosts is unable to capture individuals that may have symptomatic TB but relatively low bacterial burdens. However, as more data become available regarding the relationship between symptomatic TB and bacterial burden, future work can integrate those findings into our *HostSim* framework, perhaps by incorporating a bronchoalveolar lavage (BAL) compartment, for direct comparison to sputum samples.

In conclusion, we utilized a computational modeling framework to better understand the relationship between within-host dynamics and clinical outcomes in TB. We present *HostSim*: the first whole-host model to track events across granuloma- and host-scales. Using *HostSim*, we make predictions about relationships between immune cell counts in the blood and lungs and the role of adaptive and innate immune cells in granuloma-scale and host-scale outcomes. In particular, we predict that adaptive immunity generated in lymph nodes drives clinical classifications across hosts in TB, but that innate immunity can drive heterogeneous granuloma outcomes within a single host. We posit that *HostSim* offers a powerful computational tool that can be used in concert with experimental approaches to understand and predict events about various aspects of TB disease and therapeutics.

## Acknowledgements

This research was supported by The Wellcome Trust Delta Leap Program (DEK, JJL) and NIH Grants R01AI123093 and R01 AI50684 (DEK) and U01 HL131072 awarded to DEK and JJL. LRJ was funded by a University of Michigan Rackham Predoctoral Fellowship. Simulations also use resources of the National Energy Research Scientific Computing Center, which is supported by the Office of Science of the U.S. Department of Energy under Contract No. ACI-1053575 and the Extreme Science and Engineering Discovery Environment (XSEDE), which is supported by National Science Foundation Grant MCB140228. We thank JoAnne Flynn, Hannah Gideon and their lab members for access to previously published TB datasets.

