## Supplementary Materials for "A virtual host model of *Mycobacterium tuberculosis* infection identifies early immune events as predictive of infection outcomes"

**Supplementary Material**

*Granuloma equations*

This model, composed of a system of 20 ODEs, captures bacterial, T cell, macrophage and cytokine dynamics within a single granuloma lesion. Below, we have listed equations, variable names and a brief explanation of the dynamics of each equation. Parameter symbols generally adhere to the following guidelines: $\alpha$ parameters are growth rates. $k$ parameters are rate constants involving other variables. $\mu$ parameters are death or decay rates. $c$ parameters are half-saturation values. $Sr$ parameters are recruitment rate values to represent recruitment of cells from other areas of the body, for example. $\beta$ and $f$ parameters are scaling constants.

Extracellular bacteria concentrations within the granuloma are represented as $B_E$ across time. Extracellular bacteria can grow ($alpha20$), or can be released when infected macrophages ($M_I$) undergo apoptosis from cytotoxic T cells ($T_C$) or TNF ($F_\alpha$). Activated macrophages ($M_A$) or resting Macrophages ($M_R$) can kill extracellular bacteria.

$$\frac{dB_E}{dt} = \alpha_{20}B_E + k_{17}NM_I\left(\frac{B_I^2}{B_I^2+N^2M_I^2}\right) + k_{14a}N_{fracc}\frac{B_I}{M_I}M_I\left(\frac{\left(\frac{T_C+w_3T_1}{M_I}\right)}{\left(\frac{T_C+w_3T_1}{M_I}\right)+c_4}\right) + k_{14b}N_{fraca}\frac{B_I}{M_I}M_I\left(\frac{F_\alpha}{F_\alpha+f_9I_{10}+s_{4b}}\right) - k_2\frac{N}{2}M_R\left(\frac{B_E}{B_E+c_9}\right) - k_{15}M_AB_E - k_{18}M_RB_E - \mu_{B_E}B_E + \mu_{M_I}N_{fracd}\frac{B_I}{M_I}M_I$$

Intracellular bacteria concentrations are represented as $B_I$ across time. $B_I$ happens in the model as resting macrophages ($M_R$) engulf extracellular bacteria at a rate of $k_2$. Intracellular bacteria can die and can also become extracellular bacteria when infected macrophages undergo apoptosis. The intracellular bacterial growth term (beginning with $\alpha_{19}$) has been slightly modified from *MultiGran* (36) so it now follows logistic growth function with a carrying capacity (N) per infected macrophage ($M_I$).

$$\frac{dB_I}{dt} = \alpha_{19}\frac{B_I}{M_I}M_I\left(1-\frac{\frac{B_I}{M_I}}{N}\right) + k_2\frac{N}{2}M_R\left(\frac{B_E}{B_E+c_9}\right) - k_{17}NM_I\left(\frac{B_I^2}{B_I^2+N^2M_I^2}\right) - k_{14a}\frac{B_I}{M_I}M_I\left(\frac{\left(\frac{T_C+w_3T_1}{M_I}\right)}{\left(\frac{T_C+w_3T_1}{M_I}\right)+c_4}\right) - k_{14b}\frac{B_I}{M_I}M_I\left(\frac{F_\alpha}{F_\alpha+f_9I_{10}+s_{4b}}\right) - k_{52}\frac{B_I}{M_I}M_I\left(\frac{\left(\frac{\frac{T_C}{B_i+1}\left(\frac{T_1}{T_1+c_{T_1}}\right)+w_1T_1}{M_I}\right)}{\left(\frac{\frac{T_C}{B_i+1}\left(\frac{T_1}{T_1+c_{T_1}}\right)+w_1T_1}{M_I}\right)+c_{52}}\right) - \mu_{B_I}B_I + \mu_{M_I}\frac{B_I}{M_I}M_I$$

Resting macrophages ($M_R$) are recruited to the granuloma according to the number of activated Macrophages ($M_A$), the number of infected macrophages ($M_I$) and the concentration of TNF ($F_\alpha$) in the granuloma. $M_R$ can become activated or infected macrophages, or die.

$$\frac{dM_R}{dt} = Sr_M + \alpha_{4a}(M_A + w_2 M_I) + Sr_{4b}\left(\frac{F_\alpha}{F_\alpha + f_8 I_{10} + s_{4b}}\right) - k_2 M_R\left(\frac{B_E}{B_E + c_9}\right) - k_3 M_R\left(\frac{B_E + wB_I + \beta F_\alpha}{B_E + wB_I + \beta F_\alpha + c_8}\right)\left(\frac{I_\gamma}{I_\gamma + f_1 I_4 + f_7 I_{10} + s_1}\right) - \mu_{M_R} M_R$$

Infected macrophages ($M_I$) become infected when a resting macrophage engulfs extracellular bacteria ($B_E$). $M_I$ can burst when $B_I$ growth exceeds carrying capacity ($N$) and can die through TNF- or cytotoxic T cell- mediated apoptosis.

$$\frac{dM_I}{dt} = k_2 M_R\left(\frac{B_E}{B_E + c_9}\right) - k_{17} M_I\left(\frac{B_I^2}{B_I^2 + N^2 M_I^2}\right) - k_{14a} M_I\left(\frac{\left(\frac{T_C + w_3 T_1}{M_I}\right)}{\left(\frac{T_C + w_3 T_1}{M_I}\right) + c_4}\right) - k_{14b} M_I\left(\frac{F_\alpha}{F_\alpha + f_9 I_{10} + s_{4b}}\right) - k_{52} M_I\left(\frac{\left(\frac{T_C\left(\frac{T_1}{T_1 + c_{T_1}}\right) + w_1 T_1}{M_I}\right)}{\left(\frac{T_C\left(\frac{T_1}{T_1 + c_{T_1}}\right) + w_1 T_1}{M_I}\right) + c_{52}}\right) - \mu_{M_I} M_I$$

Activated macrophages ($M_A$) become activated through resting Macrophages ($M_R$) interactions with extracellular bacteria ($B_E$) and IFN-γ ($I_\gamma$) in the granuloma. $M_A$ can be de-activated by IL-10 ($I10$) cytokines or die.

$$\frac{dM_A}{dt} = k_3 M_R\left(\frac{B_E + wB_I + \beta F_\alpha}{B_E + wB_I + \beta F_\alpha + c_8}\right)\left(\frac{I_\gamma}{I_\gamma + f_1 I_4 + f_7 I_{10} + s_1}\right) - k_4 M_A\left(\frac{I_{10}}{I_{10} + s_8}\right) - \mu_{M_A} M_A$$

Primed CD4+ T cells ($T_0$) can proliferate at the site of the granuloma based on numbers of activated macrophages. Additionally, they are recruited to the site according to $M_I$, $M_A$, $F_\alpha$ concentrations in the granuloma. Differentiation of primed cells to effector states is based on cytokine concentrations across the granuloma. Primed cells can die.

$$\frac{dT_0}{dt} = \alpha_{1a}(M_A + w_2 M_I) + Sr_{1b}\left(\frac{F_\alpha}{F_\alpha + f_8 I_{10} + s_{4b2}}\right) + \alpha_2 T_0\left(\frac{M_A}{M_A + c_{15}}\right) - k_6 I_{12} T_0\left(\frac{I_\gamma}{I_\gamma + f_1 I_4 + f_7 I_{10} + s_1}\right) - k_7 T_0\left(\frac{I_4}{I_4 + f_2 I_\gamma + s_2}\right) - \mu_{T_0} T_0$$

Effector Th1 T cells ($T_1$) are recruited to the granuloma according to $M_I$, $M_A$, and $F_\alpha$. They are a differentiated T cell state originating from primed CD4+ T cells or effector memory CD8+ T cells. They can die from too much IFN-γ ($I_\gamma$).

$$\frac{dT_1}{dt} = \alpha_{3a}(M_A + w_2 M_I) + Sr_{3b}\left(\frac{F_\alpha}{F_\alpha + f_8 I_{10} + s_{4b1}}\right) + k_6 I_{12} T_0\left(\frac{I_\gamma}{I_\gamma + f_1 I_4 + f_7 I_{10} + s_1}\right) + k_{31} T_{4EM} M_I - \mu_{T_\gamma}\left(\frac{I_\gamma}{I_\gamma + c}\right) T_1 M_A - \mu_{T_1} T_1$$

Effector Th2 T cells ($T_2$) are recruited to the granuloma according to $M_I$, $M_A$, and $F_\alpha$. They are a differentiated T cell state originating from primed CD4+ T cells or effector memory CD4+ T cells.

$$\frac{dT_2}{dt} = \alpha_{3a2}(M_A + w_2 M_I) + Sr_{3b2}\left(\frac{F_\alpha}{F_\alpha + f_8 I_{10} + s_{4b1}}\right) + k_7 T_0 \left(\frac{I_4}{I_4 + f_2 I_\gamma + s_2}\right) + k_{32} T_{4EM} M_A - \mu_{T_2} T_2$$

Primed CD8+ T cells ($T80$) can proliferate at the site of the granuloma based on numbers of activated macrophages. Additionally, they are recruited to the site according to $M_I$, $M_A$, $F_\alpha$ concentrations in the granuloma. Differentiation of primed cells to effector states is based on cytokine concentrations across the granuloma. Primed cells can die.

$$\frac{dT_{80}}{dt} = \alpha_{1a}(M_A + w_2 M_I) + Sr_{1b}\left(\frac{F_\alpha}{F_\alpha + f_8 I_{10} + s_{4b2}}\right) + \alpha_2 T_{80}\left(\frac{M_A}{M_A + c_{15}}\right) - k_6 I_{12} T_{80}\left(\frac{I_\gamma}{I_\gamma + f_1 I_4 + f_7 I_{10} + s_1}\right) - \mu_{T_{80}} T_{80}$$

Effector CD8+ T cells ($T_8$) are recruited to the granuloma according to $M_I$, $M_A$, and $F_\alpha$. They are a differentiated T cell state originating from primed CD8+ T cells or effector memory CD8+ T cells and can die from IFN-γ ($I_\gamma$).

$$\frac{dT_8}{dt} = m\alpha_{3ac}(M_A + w_2 M_I) + mSr_{3bc}\left(\frac{F_\alpha}{F_\alpha + f_8 I_{10} + s_{4b1}}\right) + mk_6 I_{12} T_{80}\left(\frac{I_\gamma}{I_\gamma + f_1 I_4 + f_7 I_{10} + s_1}\right) + k_{34} T_{8EM} M_I - \mu_{T_{c\gamma}}\left(\frac{I_\gamma}{I_\gamma + c_c}\right) T_8 M_A - \mu_{T_C} T_8$$

Cytotoxic CD8+ T cells ($T_C$) are recruited to the granuloma according to $M_I$, $M_A$, and $F_\alpha$. They are a differentiated T cell state originating from primed CD8+ T cells or effector memory CD8+ T cells and can die from IFN-γ ($I_\gamma$).

$$\frac{dT_C}{dt} = m\alpha_{3ac}(M_A + w_2 M_I) + mSr_{3bc}\left(\frac{F_\alpha}{F_\alpha + f_8 I_{10} + s_{4b1}}\right) + mk_6 I_{12} T_{80}\left(\frac{I_\gamma}{I_\gamma + f_1 I_4 + f_7 I_{10} + s_1}\right) + k_{33} T_{8EM} M_I - \mu_{T_{c\gamma}}\left(\frac{I_\gamma}{I_\gamma + c_c}\right) T_C M_A - \mu_{T_C} T_C$$

CD4+ effector memory T cells ($T_{4EM}$) can differentiate at the site of infection into effector cell states based on numbers of infected or activated macrophages. Additionally, they are recruited to the site according to $F_\alpha$ concentrations in the granuloma. These cells can die at the site of infection.

$$\frac{dT_{4EM}}{dt} = Sr_{4EM}\left(\frac{F_\alpha}{F_\alpha + hs_{4EM}}\right) - k_{31} T_{4EM} M_I - k_{32} T_{4EM} M_A - \mu_{T_{4EM}} T_{4EM}$$

CD8+ effector memory T cells ($T_{8EM}$) can differentiate at the site of infection into effector cell states based on numbers of infected macrophages. Additionally, they are recruited to the site according to $F_\alpha$ concentrations in the granuloma. These cells can die at the site of infection.

$$\frac{dT_{8EM}}{dt} = Sr_{8EM}\left(\frac{F_\alpha}{F_\alpha + hs_{8EM}}\right) - k_{33} T_{8EM} M_I - k_{34} T_{8EM} M_I - \mu_{T_{8EM}} T_{8EM}$$

CD4+ non-specific T cells ($T_{4Non}$) represent a generic class of T cells that do not respond to Mtb antigens but are recruited to the site according to $F_\alpha$ concentrations in the granuloma. These cells do not perform effector functions within the granuloma and die at the site of infection.

$$\frac{dT_{4N\,on}}{dt} = Sr_{4N\,on}\left(\frac{F_\alpha}{F_\alpha + hs_{4N\,on}}\right) - \mu_{T_{4N\,on}}T_{4N\,on}$$

CD8+ non-specific T cells ($T_{8Non}$) represent a generic class of T cells that do not respond to Mtb antigens but are recruited to the site according to $F_\alpha$ concentrations in the granuloma. These cells do not perform effector functions within the granuloma and die at the site of infection.

$$\frac{dT_{8N\,on}}{dt} = Sr_{8N\,on}\left(\frac{F_\alpha}{F_\alpha + hs_{8N\,on}}\right) - \mu_{T_{8N\,on}}T_{8N\,on}$$

TNF ($F_\alpha$) is an inflammatory cytokine in the granuloma and is secreted by $M_I$, $M_A$, $T_1$, $T_C$ and $T_8$ cells. It also decays in the granuloma.

$$\frac{dF_\alpha}{dt} = \alpha_{30}M_I + \alpha_{31}M_A\left(\frac{I_\gamma + \beta_2(B_E + wB_I)}{I_\gamma + \beta_2(B_E + wB_I) + f_1I_4 + f_7I_{10} + s_{10}}\right) + \alpha_{32}T_1 + \alpha_{33}\left(\frac{T_C + T_8}{2m}\right) - \mu_{F_\alpha}F_\alpha$$

IFN-γ ($I_\gamma$) is an inflammatory cytokine in the granuloma and is secreted by $M_I$, $M_A$, $T_0$, $T_1$, $T_80$, $T_C$ and $T_8$ cells. It also decays in the granuloma.

$$\frac{dI_\gamma}{dt} = s_g\left(\frac{B_E + wB_I}{B_E + wB_I + c_{10}}\right)\left(\frac{I_{12}}{I_{12} + s_7}\right) + \alpha_{5a}T_1\left(\frac{M_A}{M_A + c_{5a}}\right) + \alpha_{5b}T_8\left(\frac{M_A}{M_A + c_{5b}}\right) + \alpha_{5c}M_I + \alpha_7T_0\left(\frac{I_{12}}{I_{12} + f_4I_{10} + s_4}\right) + \alpha_7T_{80}\left(\frac{I_{12}}{I_{12} + f_4I_{10} + s_4}\right) - \mu_{I_\gamma}I_\gamma$$

IL-12 ($I_{12}$) is secreted by $M_R$ and $M_A$ cells before decaying in the granuloma.

$$\frac{dI_{12}}{dt} = s_{12}\left(\frac{B_E + wB_I}{B_E + wB_I + c_{230}}\right) + \alpha_{23}M_R\left(\frac{B_E + wB_I}{B_E + wB_I + c_{23}}\right) + \alpha_8M_A\left(\frac{s}{s + I_{10}}\right) - \mu_{I_{12}}I_{12}$$

IL-10 ($I_{10}$) is secreted by $M_I$, $M_A$, $T_1$, $T_2$, $T_C$ and $T_8$ before decaying in the granuloma.

$$\frac{dI_{10}}{dt} = \delta_7(M_I + M_A)\left(\frac{s_6}{I_{10} + f_6I_\gamma + s_6}\right) + \alpha_{16}T_1 + \alpha_{17}T_2 + \alpha_{18}\left(\frac{T_C + T_8}{2m}\right) - \mu_{I_{10}}I_{10}$$

IL-4 ($I_4$) is secreted by $T_0$ and $T_2$ before decaying in the granuloma.

$$\frac{dI_4}{dt} = \alpha_{11}T_0 + \alpha_{12}T_2 - \mu_{I_4}I_4$$

***Blood and lymph node equations***

This two-compartment model represents the dynamics of specific and non-specific T cells in the LN and blood following antigen presentation by antigen-presenting cells (APCs) in the LN. Measure units are cell counts in the lymph compartment and cell/mm$^3$ in the blood compartment. The term α represents the volume of blood in μL and is used for scaling cells when they traffic between the blood compartment and the lymph compartment.

*Lymph Node CD4+ T cells*
Antigen presentation and priming in LN compartment is driven by the following equation:

$$\frac{dAPC}{dt} = -\mu_5 APC \tag{0.1}$$

which tracks antigen presenting cells (APCs) in the LN at any time during or after infection. If the number of APCs doesn't increase (a reinfection event would be an example of increasing the APC population), the APC number decreases following an exponential decay, at the rate $\mu_5$.
Naïve T cells (Eqn. (1.2)) represented by ($N_4^{LN}$) are recruited to the LN at a rate ($k_1$) dependent on cytokine production in the LN. Since we do not track cytokines in the LN model, we use APC also as a proxy for cytokine production (modeled as a Michaelis-Menten term in Eqn. (1.2)). Other terms included influx ( $\xi_1$) and efflux ( $\xi_2$), as well as mass action priming to precursor cells ($k_2$).

$$\frac{dN_4^{LN}}{dt} = \alpha\left(V_{primeN} + V_{Ninflux}\right) - V_{Nefflux} - V_{NdiffP} \tag{0.2}$$

$$V_{primeN} = k_1 N_4^B \left(\frac{APC}{APC + hs_1}\right)$$

$$V_{NdiffP} = k_2 N_4^{LN} APC$$

$$V_{Nefflux} = \xi_2 N_4^{LN}$$

$$V_{Ninflux} = \xi_1 N_4^B$$

Precursor CD4+ T cells ($P_4^{LN}$) (Eqn. (1.3)) are generated through priming of antigen-specific naïve T cells ($k_2$) as well as through re-activation of antigen-specific central memory T cells ($k_3$); both processes are expressed as mass action terms. Proliferation is modeled as logistic growth.

$$\frac{dP_4^{LN}}{dt} = \left(V_{NdiffP} + V_{CMdiffP}\right) + V_{prolif} - V_{PdiffE} - V_{PdiffCM} - \mu_6 P_4^{LN} \tag{0.3}$$

$$V_{CMdiffP} = k_3 CM_4^{LN} APC$$

$$V_{prolif} = k_4 P_4^{LN} \left(1 - \left(\frac{P_4^{LN}}{\rho_1}\right)\right)\left(\frac{APC}{APC + hs_4}\right)$$

$$V_{PdiffE} = k_5 P_4^{LN} \left(\frac{APC}{APC + hs_5}\right)$$
$$V_{PdiffCM} = k_6 P_4^{LN} \left(1 - \left(\frac{APC}{APC + hs_5}\right)\right)$$

A Michaelis-Menten term based on antigen stimulation (APC levels) was used to adjust proliferation ($k_4$) and differentiation rates ($k_5$ and $k_6$). The likelihood of precursor cells differentiating into effector cells is directly proportional to the amount of antigen stimulation ($k_5$). The opposite assumption was applied to the likelihood of precursor cells differentiating into central memory ($k_6$). A death term ($\mu_6$) ensured that the precursor population did not persist in the absence of infection. No precursor populations exit the lymph node.

Effector CD4+ T cells are modeled in Eqn. (1.4), as $E_4^{LN}$:

$$\frac{dE_4^{LN}}{dt} = V_{PdiffE} - V_{Eefflux} - V_{EdiffEM} \tag{0.4}$$
$$V_{Eefflux} = \xi_3 E_4^{LN}$$
$$V_{EdiffEM} = k_7 E_4^{LN}$$

Terms in the equation include efflux to the blood ( $\xi_3$), and a linear differentiation to the effector memory T cell population ($k_7$). We assumed that no effector T cells die in the LN (they can die in the blood).

Similar to naïve cells, central memory T cells (Eqn. (1.5)) are recruited to the lymph node ($k_8$) in addition to an influx rate ( $\xi_4$). Other terms include differentiation from precursor cells ($k_6$), reactivation to precursor cells ($k_3$) and efflux into the blood ($\xi_5$). Given their relatively long lifespan compared to the length of the *in-silico* simulation (i.e., 200 days at most) we do not have a death term in Eqn. (1.5), as $CM_4^{LN}$:

$$\frac{dCM_4^{LN}}{dt} = \alpha\left(V_{primeCM} + \xi_4 CM_4^B\right) + V_{PdiffCM} - V_{CMdiffP} - V_{CMefflux} \tag{0.5}$$
$$V_{primeCM} = k_8 CM_4^B \left(\frac{APC}{APC + hs_8}\right)$$
$$V_{CMinflux} = \xi_4 CM_4^B$$
$$V_{CMefflux} = \xi_5 CM_4^{LN}$$

Effector memory cell formation is described in Eqn. (1.6), as $EM_4^{LN}$. A linear term captures the differentiation of CD4+ effector T cells into CD4+ effector memory ($k_7$). The last term represented efflux to the blood ( $\xi_6$). Due to the longevity of these cells, we did not introduce a death term in the LN. Like effector T cells, effector memory T cells do not enter the LN directly from the blood.

$$\frac{dEM_4^{LN}}{dt} = V_{EdiffEM} - V_{EMefflux} \tag{0.6}$$
$$V_{EMefflux} = \xi_6 EM_4^{LN}$$

***Blood CD4+ T cells***

For the blood compartment, we track 4 different T cell antigen-specific phenotypes. The antigen-specific naïve CD4+ T cell blood population is modeled by Eqn. (1.7) ($N_4^B$). We have terms for a constant source supplied from the thymus (multiplied by the antigen-specific frequency λ, i.e. $\lambda s_{N_4}$) to track specific and non-specific cells, migration from the lymph node ($\xi_2$), extra recruitment to the lymph node ($k_1$), migration to the lymph node ($\xi_1$), and death ($\mu_8$).

$$\frac{dN_4^B}{dt} = \lambda s_{N_4} + \alpha^{-1}V_{Nefflux} - V_{primeN} - V_{Ninflux} - \mu_8 N_4^B \quad (0.7)$$

The values for $s_{N_4}$ and $\mu_8$ are chosen to maintain equilibrium in the total Naïve T cell populations (based on the initial conditions taken from the NHP blood data in previous work (48)).
Eqn. (1.8) describes effector CD4+ T cells dynamics ($E_4^B$) in the blood with two terms: migration from the lymph node ($\xi_3$) and death ($\mu_1$).

$$\frac{dE_4^B}{dt} = \alpha^{-1}V_{Eefflux} - \mu_1 E_4^B \quad (0.8)$$

Central memory cells in the blood (Eqn. (1.9)) ($CM_4^B$) migrate from ($\xi_5$) and to the lymph node ($\xi_4$). Central memory cells are not recruited to the site of infection.

$$\frac{dCM_4^B}{dt} = \alpha^{-1}V_{CMefflux} - V_{CMinflux} - V_{primeCM} \quad (0.9)$$

Effector memory cells in the blood (Eqn.(1.10)) ($EM_4^B$) are modeled by two terms: migration from the lymph node ($\xi_6$) and death ($\mu_2$). Similar to effector cells these were recruited to the site of infection.

$$\frac{dEM_4^B}{dt} = \alpha^{-1}V_{EMefflux} - \mu_2 EM_4^B \quad (0.10)$$

*Non-Mtb-specific CD4+ lymphocytes*
Our computational model similarly keeps track of non-specific T cells. However, non-Mtb-specific T cells do not respond to antigen, therefore, no priming occurs in any cell population and no precursor cells are generated. Also, since we assume neither effector nor effector memory T cells enter the lymph compartment from the blood, we do not model effector or effector memory cell populations within the LN compartment (as shown in Figure 2). The production of the non-specific effector cells was modeled as a source term in the blood compartment and was included to meet the assumption of homeostasis. The equations for non-Mtb-specific CD4+ T cells are shown below. Moreover, including non-Mtb-specific cells in the model makes model predictions more realistic due to the total cell numbers more accurately reflecting the actual numbers in blood.

*Naïve CD4+ non-Mtb-specific* ($N_{nc4}^{LN}$)

$$\frac{dN_{nc4}^{LN}}{dt} = \alpha\left(k_1 N_{nc4}^B\left(\frac{APC}{APC + hs_1}\right) + \xi_1 N_{nc4}^B\right) - \xi_2 N_{nc4}^{LN} \quad (1.11)$$

*Central Memory CD4+ non-Mtb-specific - LN* ($CM_{nc4}^{LN}$)

$$\frac{dCM_{nc4}^{LN}}{dt} = \alpha\left(k_8 CM_{nc4}^{B}\left(\frac{APC}{APC+hs_8}\right)+\xi_4 CM_{nc4}^{B}\right)-\xi_5 CM_{nc4}^{LN} \tag{1.12}$$

*Naïve CD4+ non-Mtb-specific – Blood* ($N_{nc4}^{B}$)

$$\frac{dN_{nc4}^{B}}{dt} = (1-\lambda)s_{N_4} + \alpha^{-1}\xi_2 N_{nc4}^{LN} - k_1 N_{nc4}^{B}\left(\frac{APC}{APC+hs_1}\right) - \xi_1 N_{nc4}^{B} - \mu_8 N_{nc4}^{B} \tag{1.13}$$

*Effector CD4+ non-Mtb-specific - Blood* ($E_{nc4}^{B}$)

$$\frac{dE_{nc4}^{B}}{dt} = s_{E_{nc4}} - \mu_1 E_{nc4}^{B} \tag{1.14}$$

As non Mtb-specific effector cells in the blood must be produced somewhere in the body, they are modeled as source and a death rate equal to that of their antigen-specific counterparts.

*Central Memory CD4+ non Mtb-specific – Blood* ($CM_{nc4}^{B}$)

$$\frac{dCM_{nc4}^{B}}{dt} = \alpha^{-1}\xi_5 CM_{nc4}^{LN} - \xi_4 CM_{nc4}^{B} - k_8 CM_{nc4}^{B}\left(\frac{APC}{APC+hs_8}\right) \tag{1.15}$$

*Effector Memory CD4+ non-Mtb-specific - Blood* ($EM_{nc4}^{B}$)

$$\frac{dEM_{nc4}^{B}}{dt} = s_{EM_{nc4}} - \mu_2 EM_{nc4}^{B} \tag{1.16}$$

*Lymph Node and Blood Mtb-specific T cells*

There are only slight differences in our modeling of CD8+ and CD4+ T cells. Importantly, the priming of Mtb-specific naïve CD8+ T cells is impacted by cytokines released by activated CD4+ T cells in the LN. Again, as we do not directly model cytokine expression in the LN; this is modeled indirectly by a Michaelis-Menten term that includes activated CD4+ T effector cells and a weighted term for precursor CD4+ T cells. We display these equations below:

$$\frac{dN_8^{LN}}{dt} = \alpha\left(k_{10} N_8^{B}\left(\frac{DC}{DC+hs_{10}}\right)+\xi_7 N_8^{B}\right)-\xi_8 N_8^{LN} - -k_{11} N_8^{LN} DC\left(\frac{[E_4^{LN}+W_{P_4}P_4^{LN}]}{[E_4^{LN}+W_{P_4}P_4^{LN}]+hs_{11}}\right) \tag{1.17}$$

$$\frac{dP_8^{LN}}{dt} = k_{11}N_8^{LN}DC\left(\frac{[E_4^{LN} + W_{P_4}P_4^{LN}]}{[E_4^{LN} + W_{P_4}P_4^{LN}] + hs_{11}}\right) + k_{12}CM_8^{LN}DC$$
$$+k_{13}P_8^{LN}\left(1 - \frac{P_4^{LN} + P_8^{LN}}{\rho_1}\right)\left(\frac{DC}{DC + hs_{13}}\right) - k_{14}P_8^{LN}\left(\frac{DC}{DC + hs_{14}}\right) -$$
$$-k_{15}P_8^{LN}\left(1 - \left(\frac{DC}{DC + hs_{14}}\right)\right) - \mu_7 P_8^{LN} \quad (1.18)$$

*Effector CD8+ T cells in lymph node*

$$\frac{dE_8^{LN}}{dt} = k_{14}P_8^{LN}\left(\frac{DC}{DC+hs_{14}}\right) - \xi_9 E_8^{LN} - k_{16}E_8^{LN} \quad (1.19)$$

*Central Memory CD8+ T cells in lymph node*

$$\frac{dCM_8^{LN}}{dt} = \alpha\left(k_{17}CM_8^{B}\left(\frac{DC}{DC + hs_{17}}\right) + \xi_{10}CM_8^{B}\right) +$$
$$k_{15}P_8^{LN}\left(1 - \left(\frac{DC}{DC + hs_{14}}\right)\right) - k_{12}CM_8^{LN}DC - \xi_{11}CM_8^{LN} \quad (1.20)$$

*Effector Memory CD8+ T cells in lymph node*

$$\frac{dEM_8^{LN}}{dt} = k_{16}E_8^{LN} - \xi_{12}EM_8^{LN} \quad (1.21)$$

*Naive CD8+ T cells in blood*

$$\frac{dN_8^{B}}{dt} = \lambda s_{N_8} + \alpha^{-1}\xi_8 N_8^{LN} - k_{10}N_8^{B}\left(\frac{DC}{DC + hs_{10}}\right) - \xi_7 N_8^{B} - \mu_9 N_8^{B} \quad (1.22)$$

*Effector CD8+ T cells in blood*

$$\frac{dE_8^{B}}{dt} = \alpha^{-1}\xi_9 E_8^{LN} - \mu_3 E_8^{B} \quad (1.23)$$

*Central Memory CD8+ T cells in blood*

$$\frac{dCM_8^{B}}{dt} = \alpha^{-1}\xi_{11}CM_8^{LN} - \xi_{10}CM_8^{B} - k_{17}CM_8^{B}\left(\frac{DC}{DC + hs_{17}}\right) \quad (1.24)$$

*Effector Memory CD8+ T cells in blood*

$$\frac{dEM_8^B}{dt} = \alpha^{-1}\xi_{12}EM_8^{LN} - \mu_4 EM_8^B \tag{1.25}$$

***Lymph Node and Blood non-Mtb-specific CD8+ T cells***
Modeled in an identical approach as the CD4+ T cell pools that were non-Mtb-specific.

*Non-Mtb-specific Naive CD8+ T cells in lymph node*

$$\frac{dN_{nc8}^{LN}}{dt} = \alpha\left(k_{10}N_{nc8}^B\left(\frac{DC}{DC+hs_{10}}\right)+\xi_7 N_{nc8}^B\right)-\xi_8 N_{nc8}^{LN} \tag{1.26}$$

*Non-Mtb-specific Central Memory CD8+ T cells in lymph node*

$$\frac{dCM_{nc8}^{LN}}{dt} = \alpha\left(k_{17}CM_{nc8}^B\left(\frac{DC}{DC+hs_{17}}\right)+\xi_{10}CM_{nc8}^B\right)-\xi_{11}CM_{nc8}^{LN} \tag{1.27}$$

*Non-Mtb-specific Naive CD8+ T cells in blood*

$$\frac{dN_{nc8}^B}{dt} = (1-\lambda)s_{N_4} + \alpha^{-1}\xi_8 N_{nc8}^{LN} - k_{10}N_{nc8}^B\left(\frac{DC}{DC+hs_{10}}\right)-\xi_7 N_{nc8}^B - \mu_9 N_{nc8}^B \tag{1.28}$$

*Non-Mtb-specific Effector CD8+ T cells in blood*

$$\frac{dE_{nc8}^B}{dt} = s_{E_{nc8}} - \mu_3 E_{nc8}^B \tag{1.29}$$

*Non-Mtb-specific Central Memory CD8+ T cells in blood*

$$\frac{dCM_{nc8}^B}{dt} = \alpha^{-1}\xi_{11}CM_{nc8}^{LN} - \xi_{10}CM_{nc8}^B - k_{17}CM_{nc8}^B\left(\frac{DC}{DC+hs_{17}}\right) \tag{1.30}$$

*Non-Mtb-specific Effector Memory CD8+ T cells in blood*

$$\frac{dEM_{nc8}^B}{dt} = s_{EM_{nc8}} - \mu_4 EM_{nc8}^B \tag{1.31}$$

***Supplementary Table 1. Parameter table for Granuloma Model and Lymph Node & Blood model parameters***

| Parameter Name Granuloma ODE | Units | Parameter Description | Minimum Value | Maximum Value |
|---|---|---|---|---|
| **Srm** | 1/day | MR recruitment rate | 0 | 0 |
| **alpha4a** | 1/day | Macrophage recruitment of MR | 0.7 | 1.0 |
| **beta** | 1/pg | Scaling factor of Falpha for MR activation | 8.65E+06 | 1.14E+07 |

| | | | | |
|---|---|---|---|---|
| **w** | N/A | Contribution of BI to MR activation | 0.26 | 0.36 |
| **w3** | N/A | Max contribution of Th1 to MI apoptosis | 0.2 | 0.8 |
| **w2** | N/A | Contribution of MI to MR recruitment | 0.9 | 1.2 |
| **Sr4b** | 1/day | Falpha dependent recruitment of MR | 592 | 864 |
| **f8** | N/A | Ratio adjustment I10/Falpha on MR recruitment | 1.74E-03 | 2.25E-03 |
| **f9** | N/A | Ratio adjustment Falpha/I10 | 0.523 | 0.673 |
| **s4b** | pg/ml | Half saturation of Falpha on MR recruitment | 2920 | 5250 |
| **k4** | 1/day | MA deactivation by I10 | 0.08 | 0.17 |
| **s8** | pg/ml | Half saturation of I10 on MA deactivation | 244 | 1003 |
| **k2** | 1/day | MR infection rate | 0.84 | 2.31 |
| **c9** | count | Half saturation of BE on MR infection | 1622 | 7868 |
| **k3** | 1/day | MR activation rate | 0.034 | 0.045 |
| **f1** | N/A | Adjustment I4/IGamma | 126 | 165 |
| **s1** | pg/ml | Half saturation of IGamma dependent MR activation | 83 | 479 |
| **c8** | count | Half saturation of BE and BI on MR activation | 1.64E+05 | 4.09E+05 |
| **nuMR** | 1/day | MR death rate | 0.004 | 0.006 |
| **k17** | 1/day | Max rate of MI bursting | 0.088 | 0.238 |
| **N** | count | Carrying capacity of MI | 5 | 25 |
| **k14a** | 1/day | T cell induced apoptosis of MI | 0.07 | 1.7 |
| **c4** | count | Half saturation of Th1/MI ratio on MI apoptosis | 397 | 951 |
| **k14b** | 1/day | Falpha induced apoptosis of MI | 0.59 | 0.92 |
| **k52** | 1/day | Cytotoxic killing of MI | 0.54 | 0.78 |
| **w1** | N/A | Max contribution of Th1 to cytotoxic killing | 0.22 | 0.74 |
| **c52** | count | Half saturation of TC on MI killing | 1.08E+05 | 2.51E+05 |
| **cT1** | count | Half saturation of Th1 on cytotoxic killing | 30 | 40 |
| **nuMI** | 1/day | MI death rate | 0.003 | 0.004 |
| **nuMA** | 1/day | MA death rate | 0.15 | 0.20 |
| **alpha1a** | 1/day | Macrophage recruitment of T0 | 0.08 | 0.57 |
| **Sr1b** | 1/day | F/alpha dependent T0 recruitment | 25007 | 54088 |
| **s4b2** | pg/ml | Half saturation of Falpha dependent T0 recruitment | 4834 | 10175 |
| **alpha2** | 1/day | Max growth rate of T0 | 0.1 | 1 |
| **c15** | count | Half saturation of MA proliferation of T0 | 5 | 25 |
| **k6** | 1/day | Max T0 to Th1 rate | 0.10 | 0.23 |
| **f7** | N/A | Effect of I10 on IGamma induced differentiation of T0 to Th1 | 8.2 | 32.3 |
| **k7** | 1/day | Max T0 to Th2 rate | 0.24 | 0.61 |
| **f2** | N/A | Adjustment IGamma/I4 | 0.2 | 0.4 |
| **s2** | pg/ml | Half saturation I4 | 429 | 955 |

| | | | | |
|---|---|---|---|---|
| **nuT0** | 1/day | T0 death rate | 0.19 | 0.25 |
| **m** | N/A | Percentage overlap between TC and T8 | 0.68 | 0.90 |
| **alpha3a** | 1/day | Macrophage recruitment of Th1 | 0.39 | 0.82 |
| **Sr3b** | 1/day | Falpha dependent recruitment of Th2 | 19 | 83 |
| **s4b1** | pg/ml | Half saturation of Falpha dependent Th1 recruitment | 6131 | 10162 |
| **alpha3a2** | 1/day | Macrophage recruitment of Th2 | 0.25 | 0.79 |
| **Sr3b2** | 1/day | Falpha dependent recruitment of Th2 | 47.5 | 99.0 |
| **nuTg** | 1/day | Igamma induced apoptosis of Th1 | 0.290 | 0.762 |
| **c** | pg/ml | Half saturation Igamma on Th1 apoptosis | 284 | 727 |
| **nuT1** | 1/day | Th1 death rate | 0.28 | 0.37 |
| **nuT2** | 1/day | Th2 death rate | 0.29 | 0.37 |
| **alpha3ac** | 1/day | Macrophage recruitment of TC and T8 | 0.25 | 0.80 |
| **Sr3bc** | 1/day | Falpha dependent recruitment of TC and T8 | 13 | 28 |
| **nuTCg** | 1/day | Igamma induced apoptosis of Tc and T8 | 0.46 | 0.90 |
| **cc** | pg/ml | Half saturation IFN-g on TC and T8 apoptosis | 337 | 673 |
| **nuTC** | 1/day | TC death rate | 0.26 | 0.33 |
| **alpha30** | pg/(ml* day) | Falpha production by MI | 0.05 | 0.10 |
| **alpha31** | pg/(ml* day) | Falpha production by MA | 0.19 | 0.82 |
| **beta2** | 1/pg | Scaling factor of Mtb for Falpha production by MA | 10466 | 13466 |
| **s10** | pg/ml | Half saturation of Igamma on Falpha production by MA | 103 | 313 |
| **alpha32** | pg/(ml* day) | Falpha production by Th1 | 0.20 | 0.34 |
| **alpha33** | pg/(ml* day) | Falpha production by T8 | 0.18 | 0.33 |
| **nuTNF** | 1/day | Falpha decay rate | 0.93 | 1.21 |
| **sg** | pg/(ml* day) | Igamma production by dendritic cells (DCs) | 2667 | 7944 |
| **c10** | count | Half saturation of Mtb on Igamma production by DCs | 980813 | 6876625 |
| **s7** | pg/ml | Half saturation of I12 on Igamma production by DCs | 538 | 883 |
| **alpha5a** | pg/day | Igamma production by Th1 | 0.54 | 0.87 |
| **c5a** | count | Half saturation of MA on Igamma production by Th1 | 304 | 687 |
| **alpha5b** | pg/day | Igamma production by T8 | 0.18 | 0.60 |
| **alpha5c** | pg/day | Igamma production by MI | 0.12 | 0.36 |
| **c5b** | count | Half saturation of MA on Igamma production by T8 | 235.8 | 846.6 |
| **alpha7** | pg/day | Igamma production by T0 | 0.04 | 0.17 |
| **f4** | N/A | Adjustment of I10/I12 on Igamma | 1.30 | 1.67 |
| **s4** | pg/ml | Half saturation of I12 on Igamma | 285 | 810 |

| | | | | |
|---|---|---|---|---|
| **nuIG** | 1/day | Igamma decay rate | 5.448 | 9.693 |
| **alpha23** | pg/day | I12 production by MR | 0.003 | 0.005 |
| **c23** | pg/ml | Half saturation of Mtb on I12 production by MR | 157 | 525 |
| **alpha8** | pg/day | I12 production by MA | 0.38 | 0.86 |
| **s12** | pg/day | Dendritic cell production of I12 | 2361 | 4061 |
| **c230** | count | Half saturation of Mtb on I12 production by DCs | 366 | 762 |
| **nuI12** | 1/day | I12 decay rate | 0.93 | 1.24 |
| **s** | pg/ml | I10 effect on I12 production by MA | 192 | 694 |
| **delta7** | pg/day | I10 production by MA | 0.38 | 0.85 |
| **s6** | pg/ml | Half saturation of I10 self | 587 | 859 |
| **f6** | N/A | Adjustment Igamma on I10 | 0.30 | 0.39 |
| **alpha16** | pg/day | I10 production by Th1 | 0.33 | 0.79 |
| **alpha17** | pg/day | I10 production by Th2 | 0.28 | 0.53 |
| **alpha18** | pg/day | I10 production by TC and T8 | 0.46 | 0.78 |
| **nuI10** | 1/day | I10 decay rate | 1.80 | 4.53 |
| **alpha11** | pg/day | I4 production by T0 | 0.01 | 0.06 |
| **alpha12** | pg/day | I4 production by Th2 | 0.02 | 0.06 |
| **nuI4** | 1/day | I4 decay rate | 2.37 | 3.09 |
| **alpha19** | 1/day | BI growth rate | 0.82 | 1.36 |
| **alpha20** | 1/day | BE growth rate | 0.25 | 0.43 |
| **Nfracc** | N/A | Fraction BI released by T cell apoptosis of MI | 0.05 | 0.07 |
| **Nfraca** | N/A | Fraction BI released by TNF apoptosis of MI | 0.05 | 0.07 |
| **k15** | 1/day | BE killing by MA | 0.0003 | 0.0011 |
| **k18** | 1/day | BE killing by MR | 0.0003 | 0.0008 |
| **Nfracd** | N/A | fraction of BI released during MI natural death to become BE | 0.0009 | 0.0011 |
| **power** | N/A | scaling factor | 2 | 2 |
| **nI** | 1/day | BI death rate | 5.95E-05 | 9.04E-05 |
| **nE** | 1/day | BE death rate | 4.11E-09 | 7.41E-09 |
| **Sr4Non** | 1/day | TNFalpha dependent recruitment of CD4 non-specific T cells | 156 | 504 |
| **hs4Non** | pg/ml | half sat of TNFalpha dependent recruitment of CD4 non-specific T cells | 5 | 50 |
| **mui4Non** | 1/day | death rate of CD4 non-specific T cells | 0.3 | 0.4 |
| **Sr8Non** | 1/day | TNFalpha dependent recruitment of CD8 non | 153 | 509 |
| **hs8Non** | pg/day | half sat of TNFalpha dependent recruitment of CD8 non | 5 | 50 |
| **mui8Non** | 1/day | death rate of CD8 non-specific T cells | 0.3 | 0.4 |
| **Sr4EM** | 1/day | TNFalpha dependent recruitment of CD4 EM T cells | 48 | 102 |
| **hs4EM** | pg/ml | half sat of TNFalpha dependent recruitment of CD4 EM T cells | 0.001 | 0.001 |

| | | | | |
|---|---|---|---|---|
| **mui4EM** | 1/day | death rate of CD4 EM T cells | 0.16 | 0.32 |
| **k31** | 1/day | differentiaton rate of CD4 EM T cells to Th1 cells | 0.04 | 0.11 |
| **k32** | 1/day | differentiaton rate of CD4 EM T cells to Th2 cells | 0.07 | 0.11 |
| **Sr8EM** | 1/day | TNFalpha dependent recruitment of CD8 EM T cells | 53 | 107 |
| **hs8EM** | pg/ml | half sat of TNFalpha dependent recruitment of CD8 EM T cells | 0.001 | 0.001 |
| **mui8EM** | 1/day | death rate of CD8 EM T cells | 0.17 | 0.31 |
| **k33** | 1/day | differentiaton rate of CD8 EM T cells to cytotoxic CD8 T cells | 0.05 | 0.11 |
| **k34** | 1/day | differentiaton rate of CD8 EM T cells to Effector CD8 T cells | 0.05 | 0.11 |
| **k99** | 1/day | killing rate constant of BI by TRM | 0.3 | 0.8 |
| **APCtimeStart** | day | when APCs leave and enter the lymph node | 5 | 28 |
| **APCtimeEnd** | day | APCs stop leaving granuloma | 50 | 105 |
| **APCleave** | N/A | Percentage of infected macs considered APCs in granuloma | 5 | 25 |
| **localDissemCFUHalf** | count | half sat CFU for local dissemination events | 6.98E+03 | 9.71E+03 |
| **localDissemLambda** | N/A | max probability of local dissemination | 0.0005 | 0.025 |
| **nonLocalDissemCFUHalf** | count | half sat CFU for non local dissemination | 5.22E+03 | 1.06E+04 |
| **nonLocalDissemLambda** | N/A | max probability of non local dissemination | 0.0001 | 0.005 |
| **MR (initial condition)** | count | Resting macrophages | 0 | 0 |
| **MI (initial condition)** | count | Infected macrophages | 1 | 1 |
| **MA (initial condition)** | count | Activated macrophages | 0 | 0 |
| **T0 (initial condition)** | count | Primed CD4+ T cells | 0 | 0 |
| **T1 (initial condition)** | count | Th1 cells | 0 | 0 |
| **T2 (initial condition)** | count | Th2 cells | 0 | 0 |
| **T80 (initial condition)** | count | Primed CD8+ T cells | 0 | 0 |
| **TC (initial condition)** | count | Cytotoxic T cells | 0 | 0 |
| **T8 (initial condition)** | count | Effector CD8+ T cells | 0 | 0 |
| **TNF (initial condition)** | pg/ml | Tumour Necrosis Factor | 0 | 0 |
| **IFNG (initial condition)** | pg/ml | interferon | 0 | 0 |
| **IL12 (initial condition)** | pg/ml | Interleukin | 0 | 0 |
| **IL10 (initial condition)** | pg/ml | Interleukin | 0 | 0 |
| **IL4 (initial condition)** | pg/ml | Interleukin | 0 | 0 |
| **BI (initial condition)** | count | Intracellular Bacteria | 1 | 1 |
| **BE (initial condition)** | count | Extracellular Bacteria | 0 | 0 |
| **CD4Non (initial condition)** | count | Nonspecific CD4 T cells | 0 | 0 |
| **CD8Non (initial condition)** | count | Nonspecific CD8 T cells | 0 | 0 |

| | | | | |
|---|---|---|---|---|
| **EMCD4** (initial condition) | count | Effector Memory CD4 T cells | 0 | 0 |
| **EMCD8** (initial condition) | count | Effector Memory CD8 T cells | 0 | 0 |
| **Parameter Name (LN & Blood)** | **Units** | **Parameter Description** | **Minimum Value** | **Maximum Value** |
| **alfa** | uL | Conversion from Blood to LN | 360000 | 360000 |
| **host_Ln** | count | Involved Lymph Nodes in Host | 5 | 5 |
| **lambda** | count | frequency of specific Naive T cells in system | 0.0001 | 0.0001 |
| **hs1** | count | half sat of Naive CD4+ T cell recruitment | 14 | 71 |
| **hs10** | count | half sat of Naive CD8+ T cell recruitment | 46 | 88 |
| **hs11** | count | half sat of Naive CD8+ T cell priming | 13 | 48 |
| **hs13** | count | half sat of precursor CD8+ T cell proliferation | 2684 | 4056 |
| **hs14** | count | half sat of precursor CD8+ T cell differentiation | 1904 | 4144 |
| **hs17** | count | half sat of Central Memory CD8+ T cell recruitment | 66 | 403 |
| **hs4** | count | half sat of Precursor CD4+ T cell proliferation | 1319 | 4318 |
| **hs5** | count | half sat of Precursor CD4+ T cell differentiation | 1257 | 3719 |
| **hs8** | count | half sat of Central Memory CD4+ T cell recruitment | 40 | 57 |
| **k1** | 1/day | Naive CD4+ T cell recruitment rate | 0.12 | 0.47 |
| **k10** | 1/day | Naive CD8+ T cell recruitment rate | 0.77 | 0.97 |
| **k11** | 1/day | Naive CD8+ T cell priming rate | 0.00010 | 0.00023 |
| **k12** | 1/day | Central Memory CD8+ T cell reactivation rate | 0.00012 | 0.00075 |
| **k13** | 1/day | Precursor CD8+ T cell proliferation rate | 0.20 | 0.80 |
| **k14** | 1/day | Precursor CD8+ T cell differentiation to Effector rate | 0.25 | 0.74 |
| **k15** | 1/day | Precursor CD8+ T cell differentiation to CM rate | 0.53 | 0.86 |
| **k16** | 1/day | Precursor CD8+ T cell differentiation to EM rate | 0.16 | 0.82 |
| **k17** | 1/day | Central Memory CD8+ T cell recruitment rate | 0.35 | 0.90 |
| **k2** | 1/day | Naive CD4+ T cell priming rate | 0.33 | 0.87 |
| **k3** | 1/day | Central Memory CD4+ T cell reactivation rate | 0.022 | 0.080 |
| **k4** | 1/day | Precursor CD4+ T cell proliferation rate | 0.30 | 1.36 |
| **k5** | 1/day | Precursor CD4+ T cell differentation to effector T cell | 0.27 | 0.90 |
| **k6** | 1/day | Precursor CD4+ T cell differentiation to central memory T cell | 0.3 | 0.9 |
| **k7** | 1/day | Effector CD4+ T cell differentiation to EM | 0.1 | 0.6 |
| **k8** | 1/day | Central Memory CD4+ T cell recruitment rate | 0.026 | 0.070 |
| **mu1** | 1/day | Effector CD4+ T cell death rate | 0.2 | 0.2 |
| **mu2** | 1/day | Effector Memory CD4+ T cell death rate | 0.04 | 0.04 |
| **mu3** | 1/day | Effector CD8+ T cell death rate | 0.2 | 0.2 |
| **mu4** | 1/day | EM CD8+ T cell death rate | 0.018 | 0.018 |
| **mu5** | 1/day | APC death rate | 0.05 | 0.05 |

| | | | | |
|---|---|---|---|---|
| **mu6** | 1/day | Precursor CD4+ T cell death rate | 0.0005 | 0.0005 |
| **mu7** | 1/day | Precurosr CD8+ T cell death rate | 0.015 | 0.015 |
| **mu8** | 1/day | Naive CD4+ T cell death rate | 0.3 | 0.3 |
| **mu9** | 1/day | Naive CD8+ T cell death rate | 0.05 | 0.05 |
| **rho1** | count | Precursor Carrying Capacity | 300000000 | 300000000 |
| **Wp4** | N/A | Weight factor for Precursor CD4+ T cell in CD8+ T cell priming | 0.7355 | 0.7355 |
| **xi11** | 1/day | Central Memory CD8+ Lymph efflux rate | 0.275 | 1.223 |
| **xi12** | 1/day | EM CD8 Lymph efflux rate | 0.219 | 1.521 |
| **xi2** | 1/day | Naive CD4+ Lymph Efflux rate | 2 | 5 |
| **xi3** | 1/day | Effector CD4+ Lymph Efflux rate | 1 | 4 |
| **xi5** | 1/day | Central Memory CD4+ Lymph Efflux rate | 1 | 4 |
| **xi6** | 1/day | Effector Memory CD4+ Lymph Efflux rate | 3 | 4 |
| **xi8** | 1/day | naive CD8 lymph efflux rate | 1 | 2 |
| **xi9** | 1/day | effector CD8 lymph efflux rate | 2 | 4 |

***Supplementary Table 2. Effect Measure comparisons for Figure 6C.***

Vargha and Delaney's A measure calculated pairwise between the three separate groups for differences between fold change of cell entry into lung. When A measure < 0.56, differences are considered small, if A measure > 0.71, the differences are considered to be large. Values are rounded to the nearest hundredth.

| | LTBI vs Active TB groups | TB eliminator vs Active TB groups | TB eliminator vs LTBI groups |
|---|---|---|---|
| CD4+ Effector T cell | 0.9 | 0.91 | 0.54 |
| CD4+ Effector Memory T cell | 0.53 | 0.64 | 0.6 |
| CD8+ Effector T cell | 0.64 | 0.84 | 0.7 |
| CD8+ Effector Memory T cell | 0.52 | 0.67 | 0.65 |

***Supplementary Table 3: PRCC values for host-scale sensitivity analysis***

| Parameter Names | Description | PRCC values |
|---|---|---|
| LN_k13 | Precursor CD8+ T cell proliferation | 0.22 |
| LN_k14 | CD8+ Precursor differentiation rate to CD8+ Effector T cell | -0.18 |

| | | |
|---|---|---|
| LN_k4 | Precursor CD4+ T cell proliferation | 0.32 |
| LN_k5 | CD4+ Precursor differentiation rate to CD4+ Effector T cell | -0.15 |

***Supplementary Table 4: PRCC values from granuloma-scale sensitivity analysis for active TB case***

| Parameter Names | Description | PRCC values |
|---|---|---|
| w3 | Max percentage contribution of Th1 cells to Fas-FasL apoptosis of MI | -0.12 |
| s4b | | 0.13 |
| k2 | MR infection rate | 0.11 |
| c9 | Half-sat of BE on MR infection | -0.14 |
| k17 | MI death rate due to BI | 0.36 |
| N | Carrying capacity of MI | 0.26 |
| k14a | Fas-FasL induced apoptosis of MI | -0.17 |
| k14b | TNF induced apoptosis of MI | -0.15 |
| Sr1b | TNF dependent recruitment of primed CD4+ T cells | -0.21 |
| s4b2 | Half-sat of TNF dependent recruitment of primed CD4+ T cells | 0.19 |
| k6 | Max rate of primed CD4+ T cells differentiating to Th1 | -0.22 |
| k7 | Max rate of primed CD4+ T cells differentiating to Th2 | 0.11 |
| s2 | Half-sat of IL-4 production | -0.1 |
| c | Half-sat of IFN-$\gamma$ on Th1 death | -0.12 |
| alpha32 | TNF produced by Th1 cells | -0.24 |
| nuTNF | Decay rate of TNF | 0.12 |
| alpha7 | IFN-$\gamma$ production by primed CD4+ T cells | -0.19 |
| s12 | IL-12 production | -0.19 |
| c230 | Half-sat of BI on IL-12 production | 0.19 |
| nuIL12 | Decay rate of IL-12 | 0.13 |
| nuIL10 | Decay rate of IL-10 | -0.13 |

| | | |
|---|---|---|
| alpha11 | IL-4 produced by primed CD4+ T cells | 0.26 |
| alpha20 | BE growth rate | 0.17 |
| k18 | BE killing by MR | -0.22 |

***Supplementary Table 5: PRCC values from granuloma-scale sensitivity analysis for TB eliminator case***

| Parameter Names | Description | PRCC values |
|---|---|---|
| k2 | MR infection rate | 0.11 |
| c9 | Half-sat of BE on MR infection | -0.14 |
| k17 | MI death rate due to BI | 0.22 |
| N | Carrying capacity of MI | 0.11 |
| k14a | Fas-FasL induced apoptosis of MI | -0.62 |
| c4 | Half-sat of cytotoxic and Th1 cells per MI on MI apoptosis | 0.17 |
| alpha11 | IL-4 produced by primed CD4+ T cells | 0.11 |
| k18 | BE killing by MR | -0.12 |

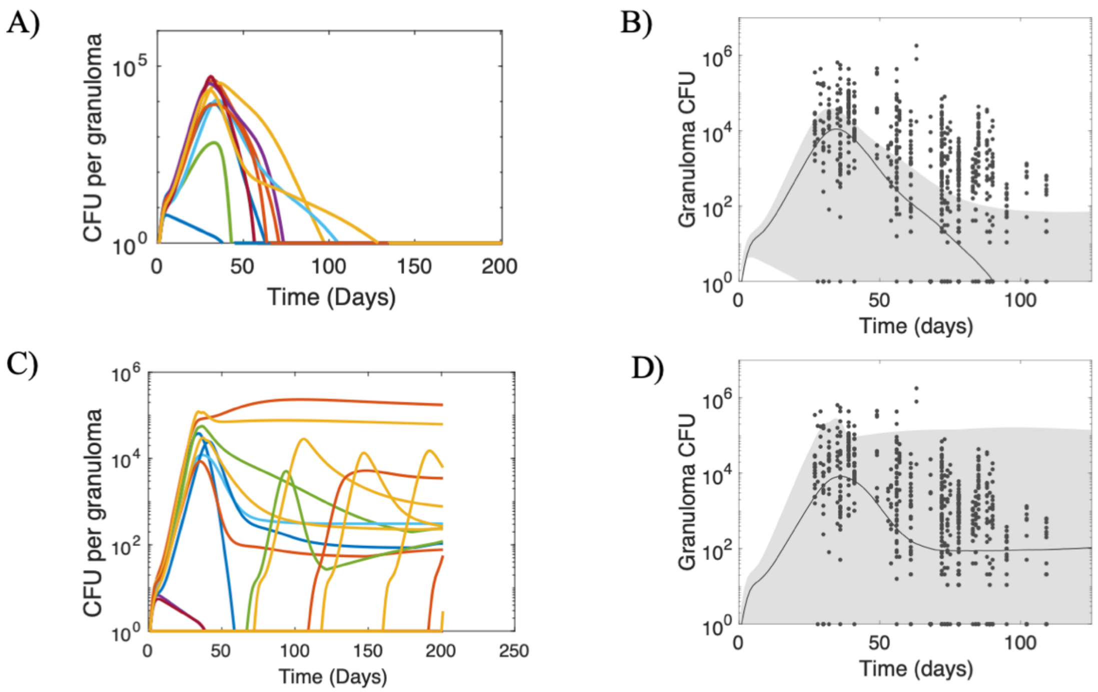


**Figure S1: Representative simulations for intra-compartment sensitivity analysis.** A) CFU trajectories within a single TB eliminator host. B) Minimum, median and maximum CFU trajectories for the TB eliminator host, re-simulated 500 times, varying only granuloma-scale parameters. C) CFU trajectories within a single active TB host. D) Minimum, median and maximum CFU trajectories for the active TB host, re-simulated 500 times, varying only granuloma-scale parameters.
